# Mitochondrial fission factor (MFF) is a critical regulator of peroxisome maturation

**DOI:** 10.1101/2020.01.08.898486

**Authors:** Josiah B. Passmore, Ruth E. Carmichael, Tina A. Schrader, Luis F. Godinho, Sacha Ferdinandusse, Celien Lismont, Yunhong Wang, Christian Hacker, Markus Islinger, Marc Fransen, David M. Richards, Peter Freisinger, Michael Schrader

## Abstract

Peroxisomes are highly dynamic subcellular compartments with important functions in lipid and ROS metabolism. Impaired peroxisomal function can lead to severe metabolic disorders with developmental defects and neurological abnormalities. Recently, a new group of disorders has been identified, characterised by defects in the membrane dynamics and division of peroxisomes rather than by loss of metabolic functions. However, the contribution of impaired peroxisome plasticity to the pathophysiology of those disorders is not well understood. Mitochondrial fission factor (MFF) is a key component of both the peroxisomal and mitochondrial division machinery. Patients with MFF deficiency present with developmental and neurological abnormalities. Peroxisomes (and mitochondria) in patient fibroblasts are highly elongated as a result of impaired organelle division. The majority of studies into MFF-deficiency have focused on mitochondrial dysfunction, but the contribution of peroxisomal alterations to the pathophysiology is largely unknown. Here, we show that MFF deficiency does not cause alterations to overall peroxisomal biochemical function. However, loss of MFF results in reduced import-competency of the peroxisomal compartment and leads to the accumulation of pre-peroxisomal membrane structures. We show that peroxisomes in MFF-deficient cells display alterations in peroxisomal redox state and intra-peroxisomal pH. Removal of elongated peroxisomes through induction of autophagic processes is not impaired. A mathematical model describing key processes involved in peroxisome dynamics sheds further light into the physical processes disturbed in MFF-deficient cells. The consequences of our findings for the pathophysiology of MFF-deficiency and related disorders with impaired peroxisome plasticity are discussed.

## 1. Introduction

Peroxisomes are highly dynamic membrane-bound organelles with key functions in cellular lipid and ROS metabolism. Defects in peroxisome biogenesis and metabolic function can result in severe disorders with developmental defects and neurological abnormalities (Dorninger et al. 2017; Wanders 2018). Peroxisome biogenesis disorders (PBDs) result from mutations in *PEX* genes, which encode proteins essential for peroxisomal membrane biogenesis and matrix protein import. PBDs, such as Zellweger Spectrum disorders, are usually characterised by a loss of functional peroxisomes. This impacts on multiple metabolic pathways (e.g., peroxisomal α- and β-oxidation of fatty acids, and the synthesis of ether-phospholipids, which are abundantly present in myelin sheaths) and results in various patient phenotypes and symptoms (Braverman et al. 2016). Peroxisomal single enzyme deficiencies (PEDs) on the other hand are caused by mutations in genes encoding a specific peroxisomal enzyme/protein and usually affect one metabolic pathway or function. The most prominent example is X-linked adrenoleukodystrophy, which is caused by mutations in the *ABCD1* gene, encoding a peroxisomal ABC transporter required for the import of very-long-chain fatty acids (VLCFAs) into the organelle (Raymond et al. 1993). In addition to PBDs and PEDs, a third group of disorders has been identified, which is characterised by defects in the membrane dynamics and division of peroxisomes rather than by loss of metabolic functions (Waterham et al. 2007; Shamseldin et al. 2012; Ebberink et al. 2012; Koch et al. 2016).

Peroxisomes can form and multiply by growth and division, a defined multistep pathway involving membrane elongation of existing peroxisomes, constriction, and membrane fission (Schrader et al. 2016). In mammals, this involves the coordinated interplay of key membrane-shaping and fission proteins such as PEX11β, FIS1, MFF, and DRP1 (encoded by the *DNML1* gene) (Schrader et al. 2016). The peroxisomal membrane protein PEX11β is involved in several steps of peroxisomal growth and division: membrane deformation to facilitate elongation (Delille et al. 2010; Opaliński et al. 2011), recruitment of the division factors MFF and FIS1 to constriction sites (Koch et al. 2005; Koch and Brocard 2012; Itoyama et al. 2013), and activation of the fission GTPase DRP1 (Williams et al. 2015). The tail-anchored membrane proteins MFF and FIS1 act as adaptor proteins for the recruitment of DRP1 to the peroxisomal membrane and interact with PEX11β (Schrader et al. 2016). With the exception of PEX11β, all proteins involved in peroxisome growth and division identified so far are also key mitochondrial division factors. FIS1 and MFF are dually targeted to both peroxisomes and mitochondria, and also recruit DRP1 to the mitochondrial outer membrane (Koch et al. 2005; Gandre-Babbe and van der Bliek 2008; Costello et al. 2017a, 2018). Mitochondria also possess the adaptor proteins MiD49 and MiD51, which are specific to mitochondria and can recruit DRP1 independent of FIS1 and MFF (Palmer et al. 2013). GDAP1 is another tail-anchored membrane protein shared by mitochondria and peroxisomes, which influences organelle fission in an MFF- and DRP1-dependent manner in neurons (Huber et al. 2013). Recently, also MIRO1, a tail-anchored membrane adaptor for the microtubule-dependent motor protein kinesin, has been shown to localise to mitochondria and peroxisomes and to contribute to peroxisomal motility and membrane dynamics (Castro et al. 2018; Okumoto et al. 2018; Covill-Cooke et al. 2020).

Patients with mutations in DRP1/DNML1, PEX11β, or MFF have been identified and often present with neurological abnormalities (Waterham et al. 2007; Shamseldin et al. 2012; Ebberink et al. 2012; Costello et al. 2018). Loss of DRP1 or MFF function leads to a block in mitochondrial and peroxisomal fission resulting in highly elongated organelles with impaired dynamics. However, the metabolic functions of both peroxisomes and mitochondria are typically not or only slightly altered, indicating that changes in organelle dynamics and plasticity are the main contributors to the pathophysiology of the disease (Waterham et al. 2007; Shamseldin et al. 2012; Koch et al. 2016; Yoon et al. 2016; Vanstone et al. 2016; Nasca et al. 2016; Gerber et al. 2017; Nasca et al. 2018; Ladds et al. 2018).

MFF deficiency displays with developmental delay, peripheral neuropathy, optic atrophy, and Leigh-like encephalopathy (Shamseldin et al. 2012; Koch et al. 2016; Nasca et al. 2018). The mitochondria in MFF-deficient patient fibroblasts show no significant alteration in oxidative phosphorylation or mtDNA (Koch et al. 2016; Nasca et al. 2018). Likewise, loss of MFF did not significantly alter the mitochondrial membrane potential, ATP levels or the redox potential of the mitochondrial matrix in neuronal cells (Lewis et al. 2018). While the majority of studies into MFF-deficiency have focused on mitochondrial dysfunction, the contribution of peroxisomal alterations to the pathophysiology is largely unknown. Similarly to DRP1 and PEX11β patients, it appears that peroxisomal metabolic function is unaltered (Koch et al. 2016; Nasca et al. 2018), with the only known peroxisome dysfunction being hyper-elongation. In this study, we assess the extent to which peroxisomal functions and properties are altered in MFF-deficient cells, giving further insight into the pathophysiological consequences of loss-of-function of MFF. We show that loss of MFF impacts on the distribution of peroxisomal marker proteins and causes the accumulation of pre-peroxisomal membrane structures. Furthermore, peroxisomes in MFF-deficient cells display alterations in peroxisomal redox state and intra-peroxisomal pH. Interestingly, elongated peroxisomes in MFF-deficient cells are not fully static, and their dynamics can be modulated, e.g. through the induction of autophagic processes. The consequences of our findings for the understanding of the pathophysiology of MFF-deficiency and related disorders with impaired peroxisome plasticity are discussed.

## 2. Materials and Methods

### 2.1. Plasmids, Antibodies and siRNAs

The plasmids and antibodies used in this study are detailed in Tables S1 and S2, respectively. PEX14 siRNA (GAACUCAAGUCCGAAAUUA) (Lee et al. 2017) and MFF siRNA (GACCAGCAGAUCUUGACCU) (Long et al. 2013) were generated by Eurofins as 21-mer siRNAs with 3’ dTdT overhangs. PEX5 siRNA (TriFECTa kit) was obtained from Integrated DNA Technologies. siGENOME Non-Targeting siRNA Control Pool (Dharmacon) and siMAX Non Specific siRNA Control 47% GC (AGGUAGUGUAAUCGCCUUG-TT, Eurofins) were used as controls.

### 2.2. Fibroblast Cell Culture and Transfection

For routine culture and morphological experiments, MFF-deficient patient skin fibroblasts and controls (Shamseldin et al. 2012; Koch et al. 2016) were cultured in Dulbecco’s Modified Eagle Medium (DMEM), high glucose (4.5 g/L) supplemented with 10% FBS, 100 U/mL penicillin and 100 μg/mL streptomycin at 37°C (5% CO_2_ and 95% humidity). The patient cells used have previously been shown to carry the following mutations in the MFF gene: c.C190T:p.Q64* (Shamseldin et al. 2012); c.184dup:p.L62Pfs*13 combined with c.C892T:p.R298* (Koch et al. 2016; patient 1); c.453_454del:p.E153Afs*5 (Koch et al. 2016; patient 2). For FRAP experiments, cells transfected with EGFP-SKL were grown on 3.5-cm glass bottom dishes (Cellview; Greiner BioOne, Germany). For assessing peroxisome degradation during starvation, cells were cultured in Hanks’ Balanced Salt Solution (HBSS) for the time indicated, and recovered in full DMEM. For assessing peroxisome alterations with microtubule depolymerisation, cells were treated with 10 μM Nocodazole (or 0.07% DMSO as a control), for four hours prior to fixation. MFF-deficient (MFF^Q64*^) and control human fibroblasts were immortalised by introduction of the SV40 large T antigen. Immortalised fibroblasts (HUFs-T) were cultured in α-modified Eagle’s medium (MEMα) supplemented with 10% FBS, 2 mM Ultraglutamine 1 (Lonza) and 1× MycoZap antibiotics (Lonza) at 37°C (5% CO_2_ and 95% humidity). Transfection of fibroblasts was performed using the Neon Transfection System (Thermo Fisher Scientific) as previously described for roGFP2 constructs (Lismont et al. 2017) and siRNA (Schrader and Schrader 2017).

### 2.2. Immunofluorescence and Immunoblotting

Unless otherwise indicated, immunofluorescence was performed 24 hours post-transfection. Cells grown on glass coverslips were fixed for 20 minutes with 4% paraformaldehyde (PFA) in PBS (pH 7.4), permeabilised with 0.2% Triton X‐100 for 10 minutes and blocked with 1% BSA for 10 minutes. Blocked cells were incubated with primary and secondary antibodies sequentially in a humid chamber for 1 hour. Cells were washed 3 times with PBS between each individual step. Finally, coverslips were washed with ddH_2_O to remove PBS and mounted on glass slides in Mowiol 4-88-containing *n*-propyl gallate as an anti-fading (Bonekamp et al. 2013).

For detection of protein levels, cells were trypsinised, washed in PBS, and centrifuged at 500×*g* for 3 min. Cell pellets were lysed and equal amounts of protein were separated by SDS-PAGE on 12.5% polyacrylamide gels. Transfer to a nitrocellulose membrane (Amersham Bioscience, Arlington Heights, IL, USA) was performed using a semi-dry apparatus (Trans-Blot SD, Bio-rad) and analysed by immunoblotting with enhanced chemiluminescence reagents (Amersham Bioscience, Arlington Heights, IL, USA).

### 2.3. Microscopy

Cell imaging was performed using an Olympus IX81 microscope with an UPlanSApo 100x/1.40 Oil objective (Olympus Optical. Hamburg, Germany). Filters sets eGFP ET (470/40 Et Bandpass filter, Beamsplitter T495 LPXR and 525/50 ET Bandpass filter [Chroma Technology GmbH, Olching, Germany]), and TxRed HC (562/40 BrightLine HC Beamsplitter HC BS 593, 624/40 BrightLine HC [Semrock, Rochester, USA]) were used. Images were taken with a CoolSNAP HQ2 CCD camera.

Live-cell imaging of roGFP2 constructs in HUFs-T fibroblasts was performed with an Olympus IX81 microscope equipped with an UPlanSApo 100x/1.40 Oil objective (Olympus Optical, Hamburg, Germany), BP390-410 and BP470-495 bandpass excitation filters, a dichromatic mirror with a cut-off at 505 nm, a BA510-550 barrier (emission) filter, and a CCD-FV2T digital black and white camera.

Confocal images of MFF^Q64*^ fibroblasts to assess peroxisomal tubule localisation with microtubules were obtained using a Zeiss LSM 880 inverted microscope, with Airyscan spatial detector array (ChA-T1 5.7, ChA-T2 6.9) for super-resolution imaging. The Alpha Plan Apochromat 100×/1.46 oil DIC M27 Elyra objective was used, with lasers 561 nm (15% power) and 488 nm (3% power).

Confocal images of the pHRed probe in fibroblasts were obtained using a Zeiss LSM 510 META inverted microscope equipped with a Plan Apochromat 63×/1.4 NA (oil/dic) objective (Carl Zeiss), using Argon excitation 458 nm and DPSS561 excitation 561 nm, with emission collection 600–620 nm. For detection of peroxisomal pHRed (pHRed-PO) the HC PL APO CS2 63×/1.4 Oil objective was used. For live‐cell imaging, cells were plated in 3.5 cm diameter glass bottom dishes (Cellview; Greiner Bio-One). MetaMorph 7 (Molecular Devices, USA) was used to adjust for contrast and brightness.

Photo-bleaching experiments were performed using a Visitron 2D FRAP system, consisting of a 405 nm/60mW diode laser. The FRAP laser was controlled by UGA-40 controller (Rapp OptoElectronic GmbH, Hamburg, Germany) and a VisiFRAP 2D FRAP control software for Meta Series 7.5.x (Visitron System, Munich, Germany) The FRAP system was coupled into a IX81 motorized inverted microscope (Olympus, Hamburg, Germany), equipped with a PlanApo 100X/1.45 Oil objective (Olympus, Hamburg, Germany). Fluorescently-labelled proteins were visualised by using a VS-LMS4 Laser-Merge-System with solid state lasers (488 nm/75mW, Visitron System, Munich, Germany). Images were captured using a Charged-Coupled Device camera (Photometric CoolSNAP HQ2, Roper Scientific, Germany). Peroxisomes in MFF-deficient fibroblasts expressing EGFP-SKL were irradiated by using 100% output power of the 405 nm laser for 150 ms with a beam diameter of 30 pixels. This was followed by immediate observation. Further details on the methods can be found in (Schuster et al. 2011a, b).

For transmission electron microscopy, fibroblast monolayers were fixed in 0.5% glutaraldehyde in 0.2 M Pipes buffer, pH 7.2, for 15 min at room temperature. Cells were then scraped from the culture dish and pelleted at 17,000 *g* for 10 min. Following three buffer washes, the cell pellet was fragmented and postfixed for 1 h in 1% osmium tetroxide (reduced with 1.5% wt/vol potassium ferrocyanide) in 0.1 M sodium cacodylate buffer, pH 7.2. Following three 5 minute washes in distilled water, the pellet fragments were dehydrated through an ethanol gradient and embedded in Durcupan resin (Sigma-Aldrich). 70-nm ultrathin sections were collected on pioloform-coated 100-mesh copper EM grids (Agar Scientific) and contrasted with lead citrate before imaging using a JEOL JEM 1400 transmission electron microscope operated at 120 kV.

### 2.4. Measurement of Peroxisomal Body Size, Tubule Size and Length, and Number

The Metamorph 7 (Molecular Devices, USA) region measurements function was used for analysis of peroxisome size in MFF-deficient fibroblasts, following calibration of distances for the magnification used. For measurement of peroxisome body and tubule width, transmission EM images were used at 80,000- and 100,000-fold magnification. For measurement of peroxisome length, immunofluorescence images were used at 100-fold magnification and the Metamorph 7 segmented line tool was used. For calculation of peroxisomal number in control fibroblasts, an in-house ImageJ (Schneider et al. 2012) macro was used, utilising the Analyze Particles function. For MFF-deficient patient fibroblasts, peroxisome number was counted manually.

### 2.5. Marker Protein Distribution Measurements

To measure the fluorescence intensity of PEX14, PMP70, catalase or EGFP-SKL over the length of a single peroxisome in fixed cells, and EGFP-SKL fluorescence following live-cell photobleaching experiments, the ImageJ (Schneider et al. 2012) Plot Profile function was used. A 2 pixel width line was drawn along the centre of the peroxisome from the body, along the tubule for a total length of 5 μm, with channels overlaid where appropriate. The fluorescence intensity for each colour channel was measured with 65 nm increments. For marker distribution measurements, data were normalised to a 0-1 scale, with 1 representing the value of the pixel with the maximum intensity of unsaturated images. For photobleaching experiments, data are presented as the mean grey value for each increment. Only peroxisomes which did not overlap with other peroxisomes were analysed.

### 2.6. Metabolic and Biochemical Analyses

Peroxisomal parameters were determined in cultured skin fibroblasts (Ferdinandusse et al. 2016). Concentrations of VLCFAs and C26:0 lysophosphatidylcholine (C26:0 lysoPC) were measured in cultured cells as described previously (Dacremont et al. 1995; Ferdinandusse et al. 2016). Peroxisomal β-oxidation of the VLCFA hexacosanoic acid (C26:0) and pristanic acid were measured as described (Wanders et al. 1995). A D3-C22:0 loading test was performed by loading cells for 3 days with deuterated (D3) C22:0 followed by fatty acid analysis with tandem mass spectrometry, essentially as previously described (Kemp et al. 2004) but with D3-C22:0 instead of D3-C24:0. Peroxisomal phytanic acid α-oxidation (Wanders and Van Roermund 1993) and the activity of dihydroxyacetone phosphate acyltransferase (DHAPAT), a key enzyme in peroxisomal ether phospholipid synthesis, were measured as described (Ofman and Wanders 1994). Immunoblot analysis was performed with cell homogenates, which were separated by SDS-PAGE and subsequently transferred onto a nitrocellulose membrane using semidry blotting. For visualisation, the secondary antibody IRDye 800 CW goat anti-rabbit was used with the Odyssey Infrared Imaging System (LI-COR Biosciences, Nebraska, USA).

### 2.7. Measurement of Subcellular Redox Dynamics

The procedures involved in the measurement of subcellular redox levels have been previously described in detail (Lismont et al. 2017). In brief, SV40 large T antigen-transformed human fibroblasts (HUFs-T) were transfected with plasmids coding for GSH/GSSG-(roGFP2) or H_2_O_2_-sensitive (roGFP2-ORP1) reporter proteins targeted to various subcellular compartments [cytosol (c-), mitochondria (mt-), or peroxisomes (po-)]. One day later, the cells were incubated for 30-60 minutes in phenol red-free culture medium and imaging was performed to visualize both the oxidized (excitation 400 nm, emission 515 nm) and reduced (excitation 480 nm, emission 515 nm) states of roGFP2. During image acquisition, the cells were maintained in a temperature-, humidity-, and CO_2_-controlled incubation chamber. For cytosolic measurements, the ROI was selected outside the nucleus. The Cell^M/xcellence software module (Olympus) was used to quantify the relative fluorescence intensities of roGFP2 at 400 and 480 nm excitation, giving a ratiometric response.

### 2.8. Measurement of Peroxisomal pH using pHRed

Peroxisomal pH was measured as previously described (Godinho and Schrader 2017). Briefly, MFF-deficient and control fibroblasts were transfected with plasmids coding for a cytosolic or peroxisomal pH-sensitive red fluorescent protein (pHRed-Cyto and pHRed-PO, respectively) (Godinho and Schrader 2017). Twenty four hours after transfection, cells were imaged using excitation wavelengths of 458 and 561 nm. Prior to image acquisition, a controlled temperature chamber was set‐up on the microscope stage at 37°C, as well as an objective warmer. During image acquisition, cells were kept at 37°C and in a HEPES-buffered CO_2_‐independent medium. For calibration, the cells were incubated in solutions of known pH (containing 5 μM nigericin) in a confocal stage chamber. ImageJ (Schneider et al. 2012) was used to calculate the 561/458 ratiometric response.

### 2.9. Statistical Analysis

Unless indicated otherwise, a two-tailed, unpaired *t*-test was used to determine statistical differences against the indicated group (*, P < 0.05; **, P < 0.01; ***, P < 0.001). Boxplots are presented with the bottom and top of each box representing the 25th and 75th percentile values, respectively; the horizontal line inside each box representing the median; and the horizontal lines below and above each box denoting the range. In the roGFP (Fig. 4B) and roGFP-ORP (Fig. 4D) box plots, these lines denote the standard deviation. Bar graphs are presented as mean ± SEM. In-text data are presented as mean ± SD. Analysis was performed from at least three independent experiments.

## 3. Results

### 3.1. Morphological characterisation of MFF-deficient peroxisomes

To visualize peroxisomes in different MFF-deficient patient skin fibroblasts (Shamseldin et al. 2012; Koch et al. 2016) under similar conditions, we processed the cells for immunofluorescence microscopy using an antibody against PEX14, a peroxisomal membrane protein. As previously reported, fibroblasts from all three MFF-deficient patients show highly elongated peroxisomes, whereas in controls peroxisomes showed a punctate staining pattern typical for human fibroblasts (**Fig. 1A**). Mitochondria in patient cells were also reported to be elongated (Shamseldin et al. 2012; Koch et al. 2016). In many cells peroxisomes were extremely long (> 30 μm); elongation was even more pronounced than in DRP1 patient fibroblasts, which also display tubular peroxisomes and mitochondria (Waterham et al. 2007; Nasca et al. 2016). The elongation of peroxisomes in MFF-deficient fibroblasts has been suggested to be the result of a constant lipid flow from the ER to peroxisomes via membrane contact sites, which are mediated by peroxisomal ACBD5 and ER-resident VAPB (Costello et al. 2017b). As peroxisomes cannot divide due to the loss of functional MFF, lipid transfer from the ER results in a pronounced growth/elongation of the peroxisomal membrane. Furthermore, re-introduction of MFF has been shown to restore the normal, punctate peroxisomal phenotype in MFF-deficient fibroblasts (Costello et al. 2017b). We transfected MFF-deficient fibroblasts with Myc-MFF using microporation, which allowed us to monitor peroxisome morphology at early time points (2-3 hours) after transfection and therefore capture the initial stages of MFF-mediated peroxisome division (**Suppl. Fig. S1**). Cells were processed for immunofluorescence using antibodies against the Myc-tag and PEX14. Two – three hours after transfection, MFF was observed to localise in spots on elongated peroxisomes (and elongated mitochondria) supporting a role in the assembly of the division machinery and the formation of division sites. Many MFF-expressing cells already contained short, dividing peroxisomes or fully divided, spherical peroxisomes (**Suppl. Fig. S1**).

**Figure 1.**
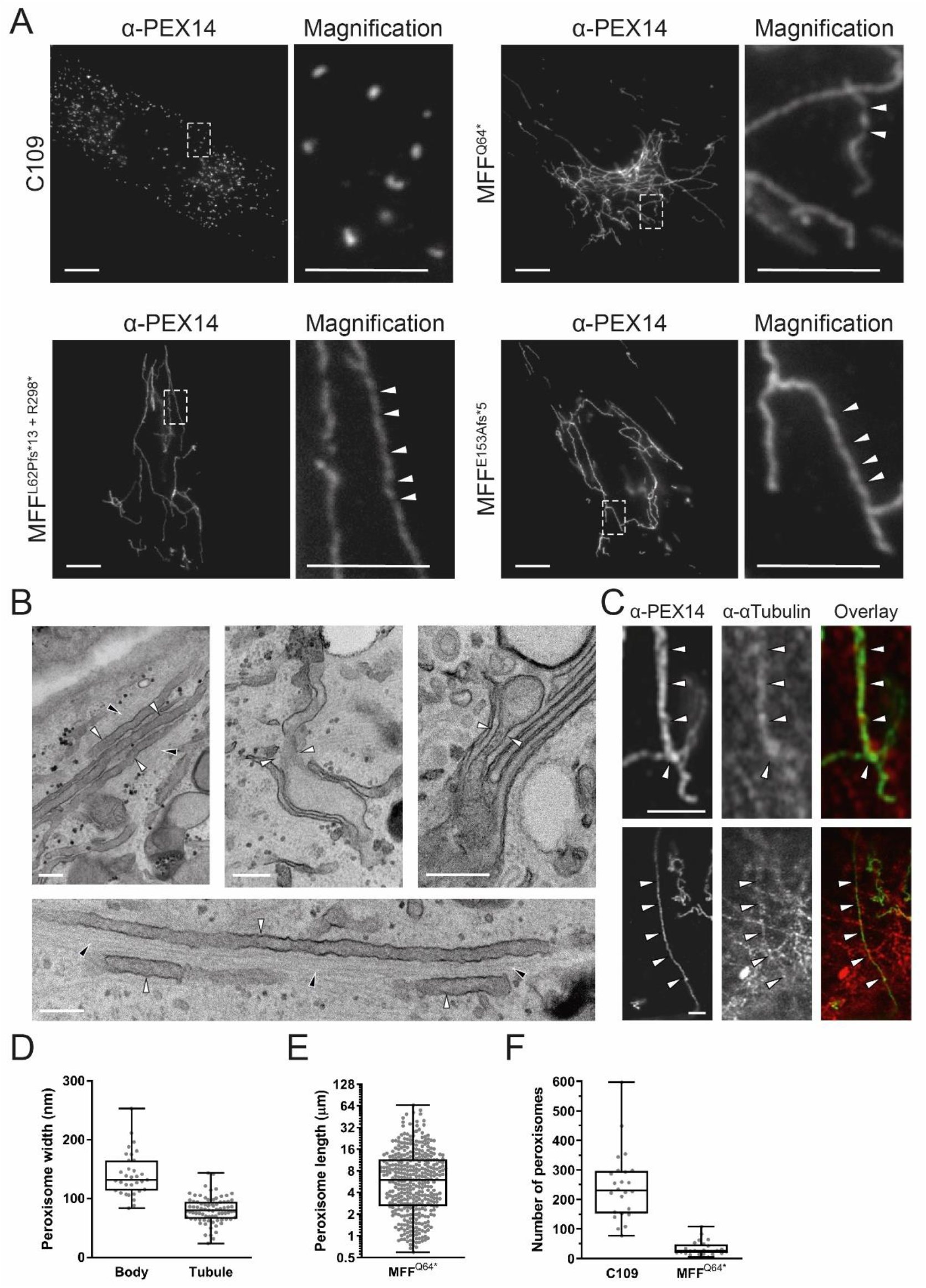
Morphological characteristics of peroxisomes in MFF-deficient patient fibroblasts are altered. (**A**) Control fibroblasts (C109) and MFF-deficient patient fibroblasts [mutations Q64* (Shamseldin et al. 2012), L62Pfs*13+R298* (Koch et al. 2016) and E153Afs*5 (Koch et al. 2016)] were processed for immunofluorescence microscopy using antibodies directed to PEX14, a peroxisomal membrane marker. Higher magnification of boxed region is shown. Arrowheads highlight potential membrane constrictions. Scale bars, 10 μm; magnification, 5 μm. (**B**) Electron micrographs of peroxisomes in MFF-deficient cells (MFF^Q64*^). White arrowheads highlight peroxisomal membrane tubules, black arrowheads indicate microtubules. Scale bars, 0.2 μm. (**C**) Confocal (Airyscan) images of peroxisomal membrane tubules (anti-PEX14) in MFF^Q64*^ cells co-stained with anti-α-tubulin. White arrowheads indicated co-localisation of peroxisomes and microtubules. Scale bars, 3 μm. (**D**) Measurement of peroxisomal width (nm) of bodies and tubules based on electron micrographs of MFF^Q64*^ fibroblasts [n = 33 (bodies), 79 (tubules)]. (**E**) Measurement of peroxisomal length (μm) from immunofluorescence images of MFF^Q64*^ patient fibroblasts (n = 392). (**F**) Quantification of peroxisome number based on immunofluorescence images of control (C109) and MFF^Q64*^ fibroblasts (n = 24). Data are from at least 3 independent experiments. ***, *p* < 0.001; two-tailed, unpaired t test.

Occasionally, peroxisomes in patient fibroblasts appeared to have a constricted, ‘beads-on-a-string’ phenotype (**Fig. 1A**, Magnifications). Such a phenotype is seen with DRP1 depletion, as peroxisomal constriction can occur independently of DRP1, but fission cannot (Koch et al. 2004). How peroxisomal constriction is mediated is still unclear. A constricted, ‘beads-on-a-string’-like peroxisome morphology in MFF-deficient cells would suggest that peroxisomal constriction can also occur independently of MFF (Ribeiro et al. 2012). However, MFF is also suggested to play a role in the constriction of the peroxisomal membrane, as it localises to peroxisomal constriction sites (Itoyama et al. 2013; Soliman et al. 2018). To confirm constricted peroxisome morphology in MFF-deficient cells, we performed electron microscopy (**Fig. 1B**). In contrast to immunofluorescence, constrictions of elongated peroxisomes were not observed in ultrastructural studies (**Fig. 1B**). Interestingly, EM revealed the presence of spherical peroxisome bodies, with a single, smaller tubule protruding from the body (**Fig. 1B**). We assume that the “constricted” appearance of peroxisomes in immunofluorescence is likely due to instability of the extremely long, delicate membrane structures during fixation with para-formaldehyde, highlighting the importance of ultrastructural studies to validate light microscopy observations. Ultrastructural studies (**Fig. 1B**) and immunofluorescence microscopy (**Fig. 1C**) show that the peroxisomal membrane tubules are frequently aligned along microtubules, which may contribute to tubule stability and maintenance.

Measurement of peroxisomes in EM micrographs revealed that peroxisome bodies are significantly larger than peroxisomal tubules (mean width, body: 141 ± 37 nm, tubule: 81 ± 22 nm) (**Fig. 1D**). The measured body width is consistent with that of spherical peroxisomes in human fibroblasts from healthy individuals typically being reported to be between 50-200 nm in width (Arias et al. 1985; Galiani et al. 2016). Peroxisome length was also quantified based on immunofluorescence data, with a wide range of lengths being present, from smaller, rod shaped peroxisomes (> 3 μm) up to very highly elongated tubules (> 30 μm) (mean length, 8.73 ± 9.2 μm) (**Fig. 1E**). As expected with a defect in division, the peroxisome number was reduced in MFF-deficient fibroblasts in contrast to controls (mean number, control fibroblasts: 244 ± 116, dMFF: 34 ± 25) (**Fig. 1F**). Overall, we reveal that peroxisomes in MFF-deficient patient fibroblasts are fewer and consist of two continuous membrane domains: a spherical peroxisome body with typical peroxisome size, and a thin, highly elongated tubular structure protruding from this body.

### 3.2. MFF deficiency does not alter standard biochemical parameters associated with peroxisomal dysfunction

Several biochemical parameters were studied to investigate peroxisomal function in cultured fibroblasts (**Table 1**). Peroxisomal α- and β-oxidation activities were measured with different radiolabelled substrates, i.e. [^14^C]-phytanic acid, pristanic acid and cerotic acid (C26:0). In addition, very long-chain fatty acid (VLCFA) metabolism was studied with a three day D3-C22 loading test, and total VLCFA levels and C26-lysophosphatidylcholine levels were determined in cell pellets (Ferdinandusse et al. 2016). No notable abnormalities were found in all three MFF-deficient cell lines providing no indication of a disturbed metabolism of VLCFAs or branched-chain fatty acids in peroxisomes. α-oxidation values were slightly higher than the reference range, but this does not indicate any dysfunction. The activity of dihydroxyacetone phosphate acyltransferase (DHAPAT), the first enzyme of the plasmalogen biosynthesis pathway located in peroxisomes, was within reference range. The intra-peroxisomal processing of the peroxisomal β-oxidation enzymes acyl-CoA oxidase 1 (ACOX1) and 3-ketoacyl-CoA thiolase was not altered, suggesting normal peroxisomal matrix protein import and processing activity in contrast to fibroblasts from a patient with a peroxisomal biogenesis disorder (**Fig. 2**). This is in line with metabolic and biochemical analyses of plasma from different MFF patients (Shamseldin et al. 2012; Koch et al. 2016; Nasca et al. 2018). We can confirm from these studies that MFF deficiency does not cause alterations to overall peroxisomal biochemical function. This is also in line with reports from other disorders affecting the dynamics and plasticity of peroxisomes (e.g. DRP1- or PEX11β-deficiency) (Waterham et al. 2007; Ebberink et al. 2012).

**Table 1.** Biochemical parameters associated with peroxisomal dysfunction are normal in MFF-deficient patient fibroblasts. Peroxisomal parameters determined in three MFF-deficient patient fibroblast cell lines MFF^L62Pfs*13+R298*^ (Koch et al. 2016), MFF^E153Afs*5^ (Koch et al. 2016), and MFF^Q64*^ (Shamseldin et al. 2012). Very long-chain fatty acid (VLCFA) levels, C26-lysophosphatidylcholine (C26-lysoPC), α- and β-oxidation activity, VLCFA metabolism (D3C22 loading test) and dihydroxyacetone phosphate acyltransferase (DHAPAT) activity were measured. A reference range of control fibroblasts from healthy individuals is shown for comparison. Data present mean of duplicate measurements. n.d., not determined; VLCFA, very long-chain fatty acid; C26-lysoPC, C26-lysophosphatdylcholine; DHAPAT, dihydroxyacetone phosphate acyltransferase.

|  | MFF<sup>L62Pfs*13+R298*</sup> | MFF<sup>E153Afs*5</sup> | MFF<sup>Q64*</sup> | Reference range |
| --- | --- | --- | --- | --- |
| **VLCFAs** (μmol/g protein) | | | | |
| C26:0 | 0.30 | 0.33 | 0.29 | 0.16-0.41 |
| C26/C22 ratio | 0.06 | 0.07 | 0.07 | 0.03-0.1 |
| **C26-lysoPC** (pmol/mg protein) | 12.3 | 9.2 | 7.0 | 2-14 |
| **Alpha-oxidation activity** (pmol/(hour.mg protein)) | 135 | 104 | n.d. | 28-95 |
| **Beta-oxidation activity** (pmol/(hour.mg protein)) |  |  |  |  |
| C26:0 | 2109 | 1505 | n.d. | 800-2040 |
| Pristanic acid | 1072 | 1099 | n.d. | 790-1072 |
| **D3C22 loading test** (μmol/g protein) | | | | |
| D3C26 (chain elongation) | 0.29 | 0.3 | 0.26 | 0.16-0.66 |
| D3C16/D3C22 ratio (beta-oxidation) | 1.25 | 1.74 | 2.27 | 0.64-2.13 |
| **DHAPAT activity** (nmol/(2hour.mg protein)) | 9.2 | 7.1 | 6.6 | 5.9-15.5 |

**Figure 2.**
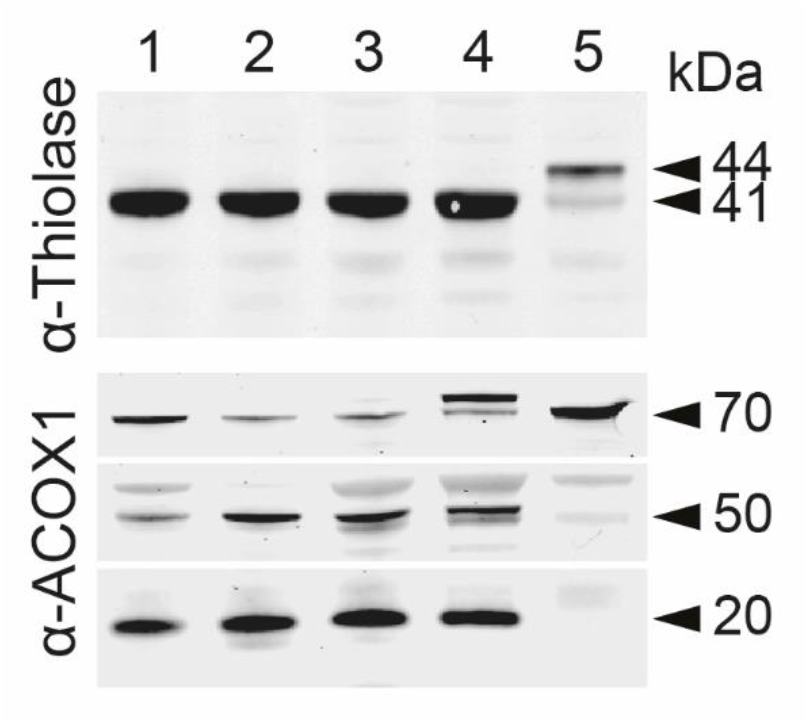
Immunoblot analysis of fibroblast homogenates from MFF-deficient patients. Antibodies were directed against peroxisomal 3-ketoacyl-CoA thiolase (upper panel) or peroxisomal acyl-CoA oxidase 1 (ACOX1; lower panel). Lanes 1-3, MFF-deficient patient fibroblasts MFF^Q64*^ (Shamseldin et al. 2012), MFF^L62Pfs*13+R298*^ (Koch et al. 2016) and MFF^E153Afs*5^ (Koch et al. 2016), respectively. Lane 4: control subject, Lane 5: fibroblasts of a patient with Zellweger Spectrum Disorder (ZSD). Results show normal proteolytic processing of 3-ketoacyl-CoA thiolase (40-kDa) and ACOX1 (50- and 20-kDa) in the MFF-deficient cell lines, whereas in the ZSD line the unprocessed bands of 3-ketoacyl-CoA thiolase (44-kDa) and ACOX1 (70-kDa) are present. Note that the protein band above the 70 kDa band of ACOX1 is non-specific.

### 3.3. Protein import into MFF-deficient peroxisomes is impaired in tubular extensions

As globular peroxisomal bodies were visible in ultrastructural studies (**Fig. 1B**) but surprisingly less visible in immunofluorescence studies with anti-PEX14, which labelled predominantly tubular structures (**Fig. 1A**), we performed co-localisation studies with anti-catalase, a prominent peroxisomal marker enzyme in the peroxisomal matrix (**Fig. 3A**). In contrast to PEX14, endogenous catalase was found to localise primarily to the spherical peroxisome bodies, with weaker fluorescence intensity along the peroxisomal tubules (**Fig. 3A**). Analysis of fluorescence intensity along single peroxisomes of both PEX14 and catalase confirmed PEX14 fluorescence primarily along tubules with some localisation in bodies, whereas catalase fluorescence was primarily detected in the peroxisomal body, with reduced intensity along the tubule (**Fig. 3A**). Peroxisomes import matrix proteins from the cytosol via dedicated import machinery at the peroxisomal membrane (Francisco et al. 2017). Matrix proteins such as catalase are imported into peroxisomes via a C-terminal peroxisomal targeting signal (PTS1). These steady-state observations imply that catalase is mainly imported into the spherical bodies, suggesting that those represent mature, import-competent structures. To test this hypothesis, we expressed a GFP-fusion protein with a C-terminal PTS1 signal SKL (GFP-SKL) in MFF-deficient cells. Cells were processed for immunofluorescence after 24 hours and labelled with anti-PEX14 antibodies (**Fig. 3B**). Similar to endogenous catalase, exogenously expressed GFP-SKL localised primarily to peroxisomal bodies, with less presence in the peroxisomal tubules (**Fig. 3B**). This was confirmed by analysis of fluorescence intensity (**Fig. 3B**). Immunofluorescence microscopy with the peroxisomal membrane markers PMP70 and PEX14 revealed co-localisation of both membrane proteins at membrane tubules (**Fig. 3C**). PMP70 also localised to the spherical bodies, where PEX14 is less prominent (**Fig. 3C**). These findings indicate that the spherical bodies represent mature, import-competent peroxisomes, whereas the tubular extensions comprise a pre-peroxisomal membrane compartment which has not yet fully acquired import competence for matrix proteins or lacks the capability to retain them. To confirm these conclusions, we performed FRAP experiments (**Suppl. Fig. S2**). Peroxisomes in MFF-deficient fibroblasts expressing GFP-SKL were photobleached followed by immediate observation through live-cell imaging. After photobleaching of the entire organelle (peroxisome body and short tubule), recovery of GFP-SKL fluorescence was first observed in the peroxisome body, indicating that recovery is due to import of GFP-SKL into the peroxisome body rather than into the tubule (**Suppl. Fig. S2**). We cannot completely exclude that there is some matrix protein import into the tubule, which may be slow or less efficient. However, our findings support our conclusion that spherical bodies are mature import competent structures, whereas the tubules represent pre-peroxisomal membrane structures which have not yet fully acquired import competence for matrix proteins or lack the capability to retain them.

**Figure 3.**
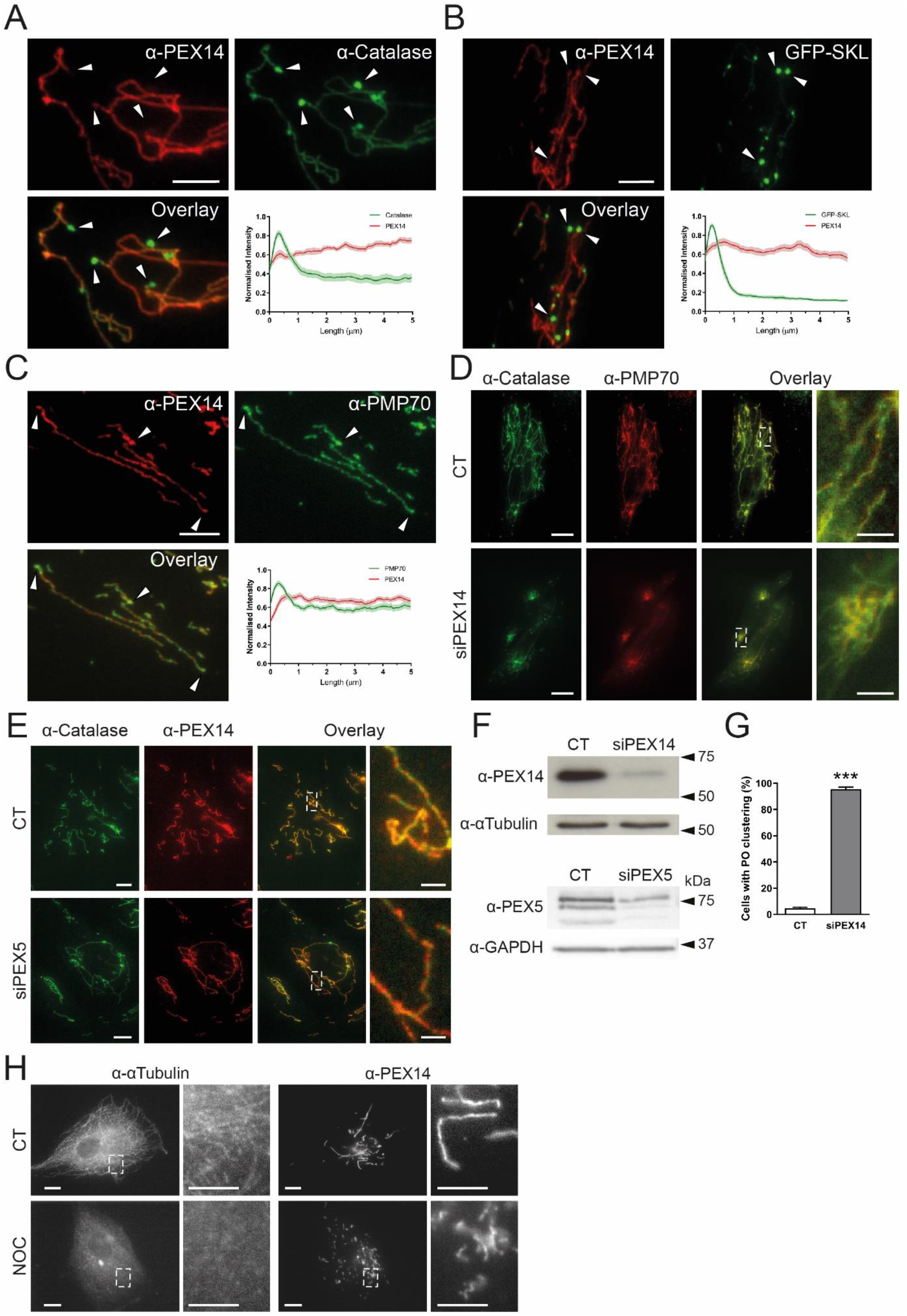
Altered marker protein distribution in MFF-deficient patient fibroblasts (MFF^Q64*^). (**A**) Patient fibroblasts were processed for immunofluorescence microscopy using antibodies against peroxisomal membrane marker PEX14 and matrix marker catalase, and fluorescence intensity measured along 5 μm of peroxisome, starting at peroxisome bodies (arrowheads) normalised to the maximum intensity. Shaded area in graphs represents the standard error of the mean (line) (n = 30). Arrowheads highlight peroxisomal bodies. Scale bar, 5 μm. (**B**) Patient fibroblasts were transfected with a plasmid encoding EGFP-SKL and processed for immunofluorescence microscopy using an antibody against PEX14. Quantification was performed as in **A**(n = 30). (**C**) As in **A**, using antibodies against membrane markers PEX14 and PMP70. Quantification was performed as in **A**, **B**(n = 30). Scale bar, 5 μm. (**D**) MFF^Q64*^ fibroblasts were transfected with control siRNA (CT) or PEX14 siRNA (siPEX14) and processed for immunofluorescence microscopy after 48 hours using antibodies against catalase and PMP70. Scale bars, 10 μm, magnification, 2 μm. (**E**) As in **D**, transfecting with control siRNA (CT), or PEX5 siRNA (siPEX5), and processing for immunofluorescence using antibodies against catalase and PEX14. Scale bars, 10 μm, magnification, 2 μm. (**F**) Immunoblotting of control (CT), PEX14 (siPEX14) or PEX5 siRNA (siPEX5) transfected patient fibroblasts, using antibodies against PEX14, PEX5 and α-tubulin or GAPDH (loading control). (**G**) Quantification of peroxisomal clustering in MFF-deficient fibroblasts either transfected with control (CT) or PEX14 siRNA (siPEX14) (n = 150). Data are from at least 3 independent experiments. ***, *p* < 0.001; two-tailed, unpaired t test. (**H**) MFF^Q64*^ patient fibroblasts were treated with 0.07% DMSO (CT), or 10 μM nocodazole (NOC) for four hours prior to processing for immunofluorescence microscopy using antibodies against α-tubulin and PEX14. Scale bars, 10 μm, magnification, 2 μm.

**Figure 4.**
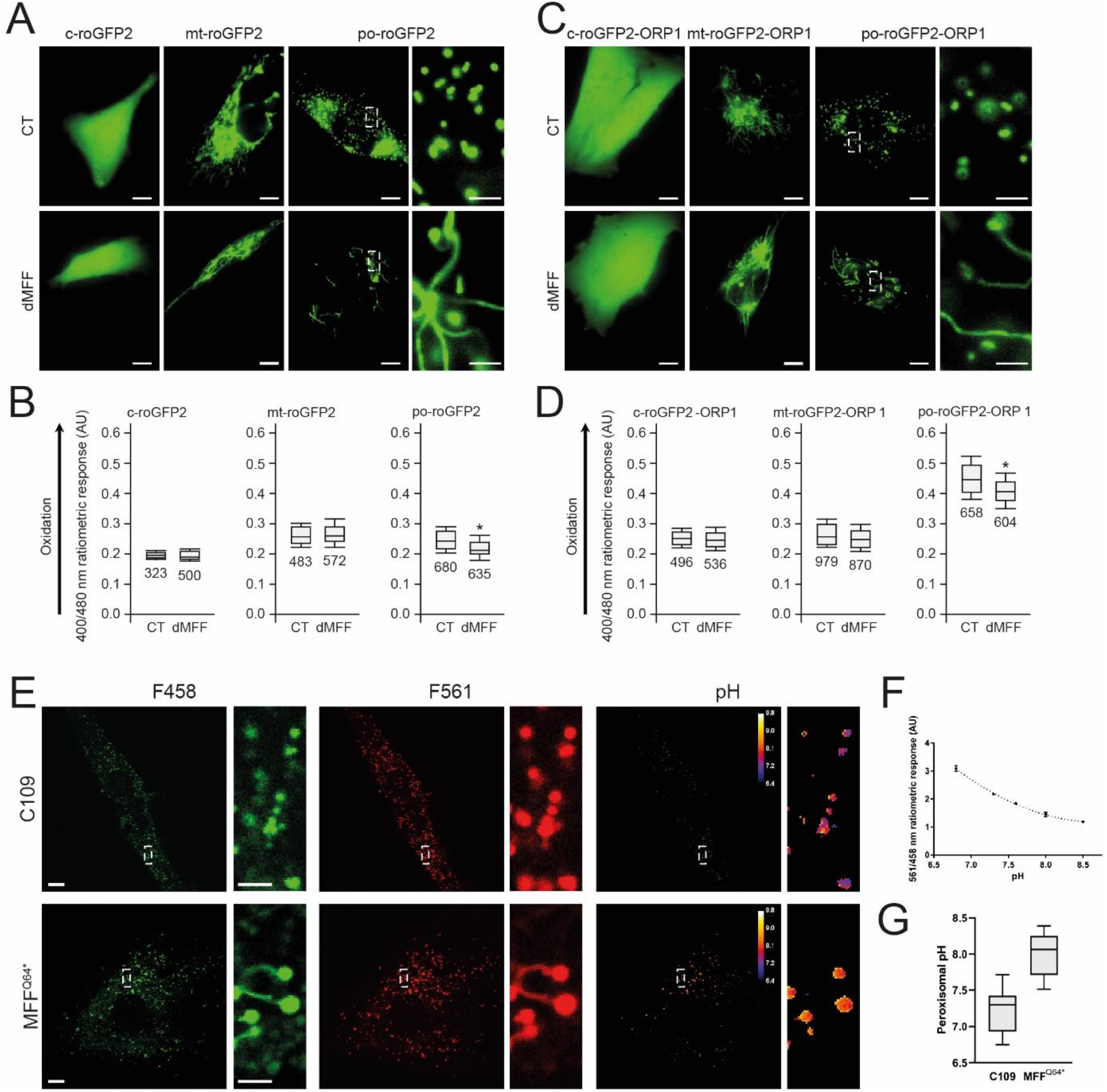
Peroxisomal redox state and pH levels are altered in MFF-deficient fibroblasts. Control (CT) or MFF-deficient (dMFF) SV40 large T antigen-transformed human fibroblasts (HUFs-T) were transfected with a plasmid encoding cytosolic (c-), mitochondrial (mt-) or peroxisomal (po-) roGFP2 (**A, B**) or roGFP2-ORP1 (**C, D**). (**A**) Distribution patterns of the respective roGFP2 proteins. Higher magnification view of po-roGFP2 is shown. (**B**) Box plot representations of the 400/480 nm fluorescence response ratios of the respective roGFP2 proteins. (**C**) Distribution patterns of the respective roGFP2-ORP1 proteins. Higher magnification view of po-roGFP2-ORP1 is shown. Note that high expression levels of the peroxisomal reporter proteins result in labelling of peroxisome tubules. (**D**) Box plot representations of the 400/480 nm fluorescence response ratios of the respective roGFP2 proteins. The bottom and top of each box represent the 25th and 75th percentile values, respectively; the horizontal line inside each box represents the median; and the horizontal lines below and above each box denote the mean minus and plus one standard deviation, respectively. The total number of measurements (two independent experiments; minimum 15 individual measurements in at least 20 randomly chosen cells) is indicated below each box plot. The data from the dMFF cell line were statistically compared with those from the CT cell line (**, p < 0.01). (**E**) Distribution patterns of pHRed-PO in control (C109) and MFF-deficient patient fibroblasts (MFF^Q64*^) at excitation wavelengths of 458 and 561 nm, along with digital visualisation of individual peroxisomal pH levels. Higher magnification views of boxed regions are indicated. (**F**) Calibration of the pHRed probe using cytosolic pHRed. The 458/561 ratiometric response is given at each pH level. AU, arbitrary units. (**G**) Quantification of peroxisomal pH in control (C109) and MFF^Q64*^ cells, converting the ratiometric response to pH using the calibration curve (n =20). Scale bars, 10 μm; magnifications, 2 μm. Data are from at least 2-3 independent experiments. *, *p* < 0.05; ***, *p* < 0.001; two-tailed, unpaired t test.

### 3.4. A role of PEX14 in maintaining peroxisomal tubule stability

As PEX14 is part of the matrix protein import machinery (Brown and Baker 2008), its predominant localisation to the peroxisomal membrane tubules (rather than the import-competent spherical bodies) is unexpected. However, additional functions for PEX14 have been suggested. Peroxisomes interact with and move along microtubules (Thiemann et al. 2000; Schrader et al. 2003; Castro et al. 2018). The N-terminal domain of PEX14 (1-78) has previously been shown to bind tubulin (Bharti et al. 2011; Theiss et al. 2012). Although PEX14 is not essential for microtubule-dependent peroxisomal motility (Castro et al. 2018), it may function as a peroxisomal microtubule docking factor. Indeed, in ultrastructural and confocal studies microtubules were frequently observed in close association with the entire length of peroxisomal tubules in MFF patient cells (**Fig. 1B, C**). Furthermore, in a previous study, we showed that highly elongated peroxisomal tubules in fibroblasts are associated with microtubules, and that tubule elongation is reduced in PEX14-deficient cells (Castro et al. 2018). Based on these observations, we hypothesised that PEX14 may be required for the stabilisation of highly elongated peroxisomal tubules. To test this, we depleted PEX14 by siRNA-mediated knock down in MFF-deficient cells (**Fig. 3D, F, G**). Peroxisomal tubules in these cells are typically stretched out in the cell, allowing for easy visualisation. However, when PEX14 was knocked down, peroxisomes lost their tubular morphology and appeared clustered or fragmented (**Fig. 3D**) (cells with clustered/fragmented morphology: control siRNA: 4.7 ± 1.2%, PEX14 siRNA: 95.3 ± 3.1%) (**Fig. 3G**). The peculiar peroxisome morphology was specific for silencing of PEX14, and was not observed after silencing of PEX5, excluding an effect of impaired peroxisomal import (**Fig. 3E**). Furthermore, peroxisome morphology was not altered after silencing of PEX11β or ACBD5 in MFF-deficient cells (Costello et al. 2017b). Clustering and fragmentation of elongated peroxisomes in MFF-deficient cells was also observed after depolymerisation of microtubules with nocodazole (**Fig. 3H**). These observations suggest a role for PEX14 in facilitating and stabilising peroxisomal membrane extensions by linking the peroxisomal membrane to microtubules. This may explain why PEX14 is predominantly localising to the highly elongated peroxisomal membranes in MFF patient cells.

### 3.5. Peroxisomal redox state and pH levels are altered in MFF-deficient fibroblasts

The metabolic parameters of peroxisomes in MFF-deficient cells were normal, in particular their different functions in lipid metabolism (**Table 1**). As peroxisomes play a role in cellular H_2_O_2_ metabolism and redox homeostasis, we also investigated these parameters (**Fig. 4**). Firstly, we assessed the glutathione disulphide (GSSG) to glutathione (GSH) ratio, a marker of oxidative balance. Therefore, MFF-deficient SV40 large T antigen-transformed human fibroblasts (HUFs-T) were transfected with a plasmid encoding cytosolic, mitochondrial or peroxisome-targeted roGFP2 (**Fig. 4A**). RoGFP2 is a highly responsive, pH-independent sensor for the glutathione redox couple, and oxidation causes a shift of its excitation maximum from 488 nm to 405 nm (Ivashchenko et al. 2011; Lismont et al. 2017). Analyses of the 400/480 ratiometric responses of peroxisome-targeted roGFP2 revealed that the intra-peroxisomal pool of glutathione is less oxidized in the MFF-deficient fibroblasts than in the control cells (**Fig. 4B**). In contrast, no alterations in the glutathione redox state could be detected in the cytosol or the mitochondrial matrix.

To monitor changes in hydrogen peroxide homeostasis, MFF-deficient HUFs-T and controls were transfected with plasmids coding for cytosolic, mitochondrial, or peroxisome-targeted roGFP2-ORP1, a H_2_O_2_-responsive variant of roGFP2 (**Fig. 4C**) (Lismont et al. 2019b). No changes in oxidation state were observed in the cytosol and mitochondria (**Fig. 4D**). However, for peroxisomes, a decreased 400/480 nm ratiometric response was seen (**Fig. 4D**), indicating reduced levels of H_2_O_2_ inside peroxisomes in MFF-deficient cells.

In addition, we used peroxisome-targeted pHRed (pHRed-PO), another ratiometric probe, to assess peroxisomal pH in MFF-deficient patient fibroblasts (Tantama et al. 2011; Godinho and Schrader 2017). Importantly, this sensor is insensitive to changes in H_2_O_2_ levels (Tantama et al. 2011). The pHRed-PO probe successfully targets to peroxisomes in control and MFF-deficient fibroblasts (**Fig. 4E**). It mainly distributes to the import-competent spherical peroxisomal bodies, but also to the membrane tubules (**Fig. 4E**). Following calibration of the pHRed probe (**Fig. 4F**), the intra-peroxisomal pH can be calculated based on the 458/561 nm ratiometric response. Interestingly, intra-peroxisomal pH in MFF-deficient fibroblasts was found to be more alkaline than in control fibroblasts (**Fig. 4G**) (mean peroxisomal pH, control: 7.24 ± 0.30, patient fibroblasts: 8.00 ± 0.29).

Overall, these findings point towards alterations in the peroxisomal redox environment. Specifically, we observed a decrease in the GSSG/GSH ratio and H_2_O_2_ levels in MFF-deficient fibroblasts. In addition, we have shown that absence of MFF results in a more alkaline intra-peroxisomal pH. This suggests that MFF-deficiency may compromise normal peroxisomal redox regulation.

### 3.6. Highly elongated peroxisomes in MFF-deficient fibroblasts can be degraded by autophagic processes

Autophagic processes are important for the maintenance of cellular homeostasis and the integrity of organelles (Anding and Baehrecke 2017). Peroxisome homeostasis is achieved via a tightly regulated interplay between peroxisome biogenesis and degradation via selective autophagy (pexophagy) (Eberhart and Kovacs 2018). It is still unclear if highly elongated peroxisomes are spared from pexophagy, e.g. due to physical limitations, as the elongated peroxisomes may not fit into the autophagosome. Such a scenario would prevent degradation of peroxisomes and could have pathophysiological consequences.

To examine if highly elongated peroxisomes in MFF-deficient fibroblasts can be degraded by autophagic processes, we first induced pexophagy by the expression of a fragment of peroxisomal biogenesis protein PEX3. Expression of the first 44 amino acids of the peroxin PEX3, which can insert into the peroxisome membrane, was observed to cause complete removal of peroxisomes (Soukupova et al. 1999). When expressing *Hs*PEX3(1-44)-EGFP in control fibroblasts (**Fig. 5A, B**), peroxisomes were greatly reduced in number, with many GFP expressing cells showing almost complete loss of PEX14 labelling (**Fig. 5A, C109**). As reported earlier, loss of peroxisomes resulted in mistargeting of HsPEX3(1-44)-EGFP to the mitochondria (Soukupova et al. 1999) (**Suppl. Fig. S3**). Interestingly, in MFF-deficient fibroblasts, expression of *Hs*PEX3(1-44)-EGFP also caused a marked reduction of peroxisomes (**Fig. 5A, middle panel, B**) or complete loss of PEX14 labelling (**Fig. 5A, lower panel, B**). Increased mitochondrial mistargeting of *Hs*PEX3(1-44)-EGFP was observed with increased loss of peroxisomes (**Fig. 5A; Suppl. Fig. S3**).

**Figure 5.**
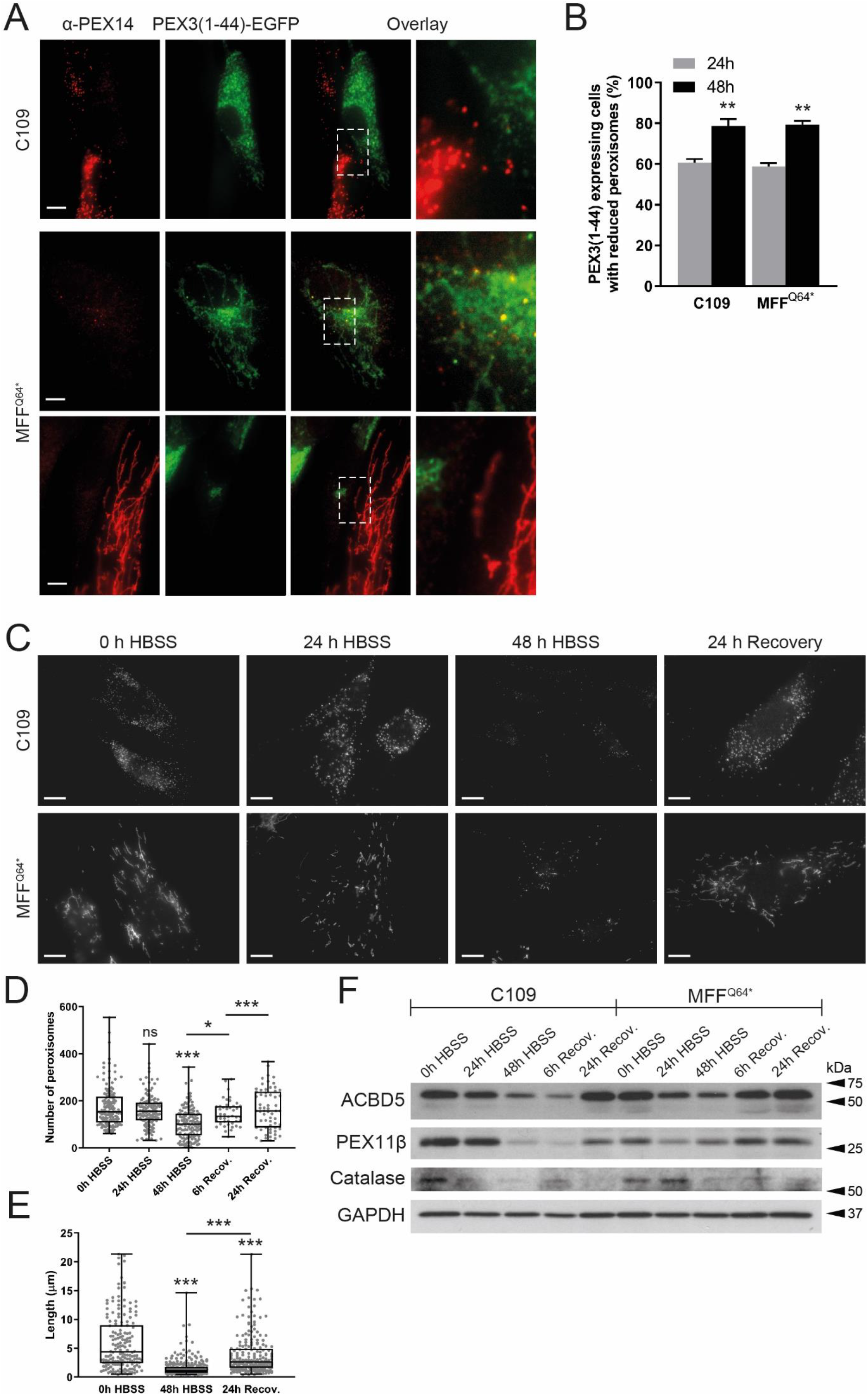
Degradation of peroxisomes in MFF-deficient patient fibroblasts. (**A**) Human control (C109) or MFF-deficient (MFF^Q64*^) fibroblasts were transfected with a plasmid coding for *Hs*PEX3(1-44)-EGFP to induce peroxisome degradation and processed for immunofluorescence after 24 and 48 hours using antibodies against PEX14. Note the almost complete loss of PEX14, and mistargeting of *Hs*PEX3(1-44)-EGFP to mitochondria when peroxisomes are lost (Soukupova et al. 1999) (**Suppl. Fig. S3**). Scale bars, 10 μm. magnification, 2 μm (**B**) Quantification of *Hs*PEX3(1-44)-EGFP expressing cells (control fibroblasts, C109; MFF-deficient, MFF^Q64*^) showing reduced peroxisomes after 24 and 48 hours (n = 150).. ***, *p* < 0.001; two-tailed, unpaired t test. (**C**) Human control (C109) and MFF-deficient fibroblasts (MFF^Q64*^) were incubated in Hanks’ Balanced Salt Solution (HBSS) to induce peroxisome degradation and processed for immunofluorescence after 0, 24 and 48 hours and after 24 hours recovery in complete culture medium using antibodies against PEX14. Scale bars, 10 μm. (**D**) Quantification of the number of peroxisomes in C109 control fibroblasts following incubation in HBSS and recovery in complete culture medium (see **C**) [n = 62 (24h Recovery) to 139 (48h HBSS)]. ***, *p* < 0.001, *, p < 0.1, ns, not significant, Ordinary one-way ANOVA with Tukey’s multiple comparisons test. (**E**) Quantification of peroxisome length in MFF^Q64*^ fibroblasts following 0, 48 hours of HBSS treatment, and after 24 hours of recovery in complete culture medium [n = 167 (0h HBSS) to 297 (48h HBSS)]. Data are from at least 3 independent experiments. ***, *p* < 0.001, Ordinary one-way ANOVA with Tukey’s multiple comparisons test. (**F**) Immunoblot of cell lysates from control (C109) and MFF-deficient fibroblasts (MFF^Q64*^) which were incubated in HBSS for 0, 24, and 48 hours, and after 6 and 24 hours of recovery in complete culture medium. Antibodies against the peroxisomal membrane proteins ACBD5, PEX11β and Catalase were applied. Anti-GAPDH was used as a loading control. Equal amounts of protein were loaded. Molecular mass markers (kDa) are indicated on the right.

To examine peroxisome degradation under more physiological conditions, we applied nutrient deprivation. Limiting amino acids in the growth media of cells has been previously shown to induce removal of peroxisomes (Sargent et al. 2016). For assessing peroxisome degradation, controls and MFF-deficient fibroblasts were cultured in Hanks’ Balanced Salt Solution (HBSS), which lacks amino acids. After 0, 24 and 48 hours, cells were processed for immunofluorescence using anti-PEX14 as a peroxisomal marker (**Fig. 5C**). In control cells, we observed a marked decrease in spherical peroxisomes, with often only a few organelles remaining after 48 hours in HBSS (**Fig. 5C, D**). As with *Hs*PEX3(1-44)-EGFP, we also observed a decrease in peroxisomes in nutrient-deprived MFF-deficient cells, which was accompanied by a significant reduction in peroxisomal length (mean peroxisomal length, 0 hours HBSS: 6.08 ± 4.90 μm, 48 hours HBSS: 1.55 ± 1.43 μm) (**Fig. 5C, E**). The reduction in peroxisomes was accompanied by a reduction in peroxisomal marker proteins (**Fig. 5F**). Peroxisomes and protein levels recovered in control and MFF-deficient cells after switching to complete culture medium for 24 hours (**Fig. 5C-F**). Interestingly, the switch to complete growth medium resulted in the recovery of elongated peroxisomes (mean peroxisomal length, 24 hours recovery: 3.84 ± 3.40 μm) (**Fig. 5E**), indicating that peroxisomes in MFF-deficient fibroblasts are still dynamic under certain conditions. Overall, these data show that highly elongated peroxisomes in MFF-deficient cells are not spared from autophagic processes and are capable of being degraded.

## 4. Discussion

Whereas dysfunctional peroxisome metabolism and associated diseases are generally well studied, the consequences and pathophysiology caused by specific disruption to peroxisome dynamics and plasticity are less clear. Mutations in DRP1, MFF or PEX11β have been linked to defects in the membrane dynamics and division of peroxisomes rather than to loss of metabolic functions (Waterham et al. 2007; Shamseldin et al. 2012; Ebberink et al. 2012; Koch et al. 2016; Taylor et al. 2017; Nasca et al. 2018). This is in contrast to the classical peroxisome biogenesis disorders (e.g. Zellweger spectrum disorders) or single enzyme deficiencies and can complicate diagnosis through metabolic biomarkers. Despite considerable progress in the field, the precise molecular functions of several of the proteins regulating peroxisomal plasticity remain to be determined as well as the contribution of impaired peroxisomal dynamics to the pathophysiology of the above disorders. In line with this, depletion of PEX11β in epidermal cells was recently reported to result in abnormal mitosis and organelle inheritance, thus affecting cell fate decisions (Asare et al. 2017). As DRP1 and MFF also localise to mitochondria, and as loss of DRP1 or MFF function also inhibits mitochondrial division, focus has so far mainly been on mitochondrial properties under those conditions. Here, we assessed the extent to which peroxisomal functions and properties are altered in MFF-deficient cells.

There are currently six patients with MFF-deficiency identified, with various mutations in the MFF protein shown; c.C190T:p.Q64* (Shamseldin et al. 2012); c.184dup:p.L62Pfs*13 combined with c.C892T:p.R298* (Koch et al. 2016); c.453_454del:p.E153Afs*5 (Koch et al. 2016); and most recently c.C892T:p.R298* alone (Nasca et al. 2018). Patient skin fibroblasts show a loss of MFF function with mitochondrial and peroxisomal hyper-elongation, and the patients themselves present with neurological abnormalities, showing developmental delay, peripheral neuropathy, optic atrophy, and Leigh-like encephalopathy (Shamseldin et al. 2012; Koch et al. 2016; Nasca et al. 2018). We confirmed a similar degree of peroxisomal hyper-elongation in skin fibroblasts from three different, previously characterized patients suffering from MFF-deficiency when maintained under the same culture conditions [c.C190T:p.Q64* (Shamseldin et al. 2012); c.184dup:p.L62Pfs*13 combined with c.C892T:p.R298* (Koch et al. 2016); c.453_454del:p.E153Afs*5 (Koch et al. 2016)]. Furthermore, peroxisomal biochemical parameters related to fatty acid α- and β-oxidation, plasmalogen biosynthesis, or matrix protein import/processing did not reveal any deficiencies in fibroblasts from those patients. This is in agreement with biochemical studies in other MFF-deficient patient fibroblasts (Koch et al. 2016; Nasca et al. 2018). Overall, these findings support the notion that defects in the membrane dynamics and division of peroxisomes rather than loss of metabolic functions contribute to the disease pathophysiology.

Similar observations in PEX11β- or DRP1-deficient cells (Waterham et al. 2007; Ebberink et al. 2012) have led to the general assumption that defects in peroxisomal dynamics and division result in elongated peroxisomes, which are, however, largely functional and otherwise normal. We now reveal in MFF-deficient cells that this is not the case. We show that the elongated peroxisomes in those cells are composed of a spherical body, which represents a mature, import-competent peroxisome, and of thin, tubular extensions, which likely represent pre-peroxisomal membrane compartments; not yet fully import-competent for peroxisomal matrix proteins. An alternative interpretation may be that the tubular structures are to some degree import-competent but lack mechanisms to retain the imported matrix proteins. Such a mechanism for retaining matrix proteins may be provided by membrane constriction, which is impaired in MFF-deficient cells.

These observations are consistent with the proposed multi-step maturation model of peroxisomal growth and division and with previous data on tubular membrane extensions after expression of PEX11β (Delille et al. 2010; Schrader et al. 2012, 2016). In this respect, elongated peroxisomes in MFF-deficient cells resemble those observed after expression of a division-incompetent PEX11β, which also results in elongated peroxisomes with an import-competent spherical body and a pre-peroxisomal membrane expansion (Delille et al. 2010). In contrast, elongated peroxisomes in DRP1-depleted cells are constricted, with a “beads-on-a string” like appearance, and the interconnected spherical peroxisomes (“beads”) are import-competent for matrix proteins (Koch et al. 2004). These constrictions may therefore provide a mechanism to retain matrix proteins. This indicates that a defect in MFF influences peroxisome division earlier than a defect in DRP1, and results in a maturation defect of elongated peroxisomes, which are unable to constrict and to subsequently import and/or retain matrix proteins. Re-expression of MFF in the MFF-deficient fibroblasts early on results in a spot-like localization of MFF on elongated peroxisomes indicating a role for MFF in the assembly of the division machinery. In line with this, it has recently been shown that MFF can act as a sensor but also potentially as an inducer of mitochondrial constriction (Helle et al. 2017). We propose that MFF deficiency, which impairs peroxisomal membrane constriction and proper assembly of the division machinery, blocks further maturation of the pre-peroxisomal membrane compartment.

This means that, although the number of fully functional peroxisomes is reduced and matrix proteins are largely restricted to the mature spherical bodies, membrane surface area and volume of the peroxisomal compartment are increased in MFF-deficient cells (mean estimated total surface area, control fibroblasts: 1.55×10^7^ ± 7.29×10^6^ nm^2^, dMFF: 1.15×10^8^ ± 6.57×10^8^ nm^2^; mean estimated total volume, control fibroblasts: 4.1×10^8^ ± 1.94×10^8^ nm^3^, dMFF 2.5×10^9^ ± 1.45×10^9^ nm^3^) (**Suppl. Fig. S4**), as well as the surface area to volume ratio (mean estimated SA:V, control fibroblasts: 0.038 ± 0.001, dMFF: 0.046 ± 0.005) (**Suppl. Fig. S4**). This likely explains why biochemical functions of elongated peroxisomes are overall normal under standard conditions. However, it can be speculated that sudden environmental changes (e.g. an increase in peroxisomal substrates via nutrients/diet or stress conditions), which require increased peroxisomal metabolic activity and number, will overwhelm the capacity of the peroxisomal compartment in MFF-deficient cells. This may also explain why mild alterations of peroxisomal metabolism are occasionally observed in patients with defects in peroxisomal dynamics and division (Waterham et al. 2007; Ebberink et al. 2012; Taylor et al. 2017). Furthermore, peroxisomes in patient cells may be less able to cope with increased expression of peroxisomal matrix enzymes or PMPs. Those may accumulate in the cytoplasm and may be degraded or mistargeted (e.g. to mitochondria) due to the reduced number of import-competent peroxisomes (Ebberink et al. 2012).

We also show that peroxisomal matrix and membrane proteins do not distribute evenly along the elongated peroxisomes in MFF-deficient cells. Endogenous catalase or exogenously expressed GFP-SKL predominantly localise to the spherical body, whereas PEX14 localises predominantly to the tubular membrane extensions. A heterogeneous distribution of peroxisomal proteins during membrane growth and division has been reported previously (Delille et al. 2010; Cepińska et al. 2011). The specific mechanisms which restrict the mobility of the peroxisomal proteins and keep them within the spherical or tubular membrane domains are still unknown, but may depend on protein oligomerization and/or a specific lipid environment. However, the prominent localisation of PEX14, a component of the docking/translocation complex for matrix protein import, to the tubular peroxisomal membranes in MFF-deficient cells is unusual. It is possible that PEX14, which has been reported to interact with microtubules (Bharti et al. 2011; Theiss et al. 2012), may also act as a peroxisome-microtubule docking factor: it predominantly localises to the peroxisomal membrane extensions in MFF patient cells and may anchor them to microtubules in order to stabilise those highly elongated, delicate membrane structures and to facilitate membrane extension. The membrane topology of PEX14 is poorly defined, but a recent study suggested that the N-terminal domain is protease-protected and may not be exposed to the cytosol (Barros-Barbosa et al. 2019). Such a topology may be inconsistent with tubulin-binding, but it is possible that different populations or complexes of PEX14 exist which may fulfil different functions at the peroxisomal membrane.

Peroxisomes are oxidative organelles with important roles in cellular redox homeostasis (Fransen and Lismont 2018). Alterations in their redox metabolism have been suggested to contribute to aging and the development of chronic diseases such as neurodegeneration, diabetes, and cancer (Fransen and Lismont 2019). Using genetically encoded fluorescent sensors with ratiometric readout in live-cell approaches, we revealed alterations in the glutathione redox potential within peroxisomes of MFF-deficient fibroblasts, which was less oxidising compared to controls. In addition, we detected reduced levels of peroxisomal H_2_O_2_ in these cells. Given that the peroxisomal parameters (**Table 1**) and catalase levels (**Fig. 5F**) are similar in control and MFF-deficient human fibroblasts, the possible mechanisms underlying these observations remain a subject of speculation. In this context, it is interesting to note that in a previous study in which mouse embryonic fibroblasts were cultured in medium containing 1,10-phenanthroline, a Zn^2+^-chelating compound that induces oxidative stress and disrupts peroxisomal and mitochondrial function (Coyle et al. 2004; Jo et al. 2015), the intra-peroxisomal redox state in tubular peroxisomal compartments was observed to be slightly lower than in spherical bodies (Lismont et al. 2017). Given that (i) peroxisome-derived H_2_O_2_ can easily cross the peroxisomal membrane (Lismont et al. 2019a), and (ii) the surface to volume ratio is larger in the tubular structures, this may be explained by the fact that H_2_O_2_ can diffuse faster out of the tubular structures than out of the spherical bodies. Alternatively, as this study indicates that matrix proteins are predominantly imported into the spherical bodies and less into the peroxisomal tubules (**Fig. 3; Suppl. Fig. S2**), the lower values for peroxisomal redox parameters in the tubular structures may also be due to the fact that these structures contain less H_2_O_2_-producing oxidases. However, in contrast to what was observed before in cells cultured in the presence of 1,10-phenanthroline, no significant differences in the glutathione redox state or H_2_O_2_ levels could be detected between the spherical and tubular structures in MFF-deficient cells (data not shown). Importantly, the glutathione redox balance and hydrogen peroxide levels in the cytosol and mitochondria were similar to controls, indicating peroxisome-specific alterations due to loss of MFF-function. Peroxisome-derived H_2_O_2_ may be an important signalling messenger that controls cellular processes by modulating protein activity through cysteine oxidation (Fransen and Lismont 2019). However, the precise interrelationship between peroxisomal redox metabolism, cell signalling, and human disease remains to be elucidated. Further insight may come from the identification of primary targets for peroxisome-derived H_2_O_2_. We also revealed changes in the peroxisomal pH in MFF-deficient fibroblasts, which was more alkaline than in controls. The pI of most peroxisomal enzymes is basic, and consistent with this, an alkaline pH has been reported for the peroxisomal lumen (Dansen et al. 2000; van Roermund et al. 2004; Godinho and Schrader 2017). Studies addressing peroxisomal pH under disease conditions are scarce, but a more acidic peroxisomal pH has been reported in fibroblasts from patients suffering from Rhizomelic Chondrodysplasia Punctata type 1, a PBD based on a defect in the import receptor PEX7 and impaired matrix protein import of PTS2-containing cargo (Dansen et al. 2000). It remains to be determined if those changes are the result of slightly altered metabolic activity and/or changes in membrane properties which impact on peroxisomal membrane channels/transporters. In line with this, calcium influx into peroxisomes has been reported to induce a minor increase of peroxisomal pH (Lasorsa et al. 2008). Whether peroxisomes possess a proton pump is still debated, but it has been suggested that a peroxisomal proton gradient may be needed to drive other transport processes across the peroxisomal membrane (Rottensteiner and Theodoulou 2006).

It is suggested that a block in peroxisome fission (e.g., due to mutations in MFF or DRP1), which results in the formation of larger, elongated organelles, may have deleterious effects on the mobility of peroxisomes, on synaptic homeostasis, and pexophagy (Schrader et al. 2014). We show here that highly elongated peroxisomes in MFF-deficient fibroblasts can be degraded by autophagic processes, which were induced by expression of a fragment of PEX3 [*Hs*PEX3(1-44)] (Soukupova et al. 1999) or by amino acid starvation. Highly elongated mitochondria, for example, were reported to be spared from mitophagy under starvation conditions (Rambold et al. 2011; Gomes et al. 2011). Our data reveal that elongated peroxisomes are not spared from autophagic processes, e.g. due to physical limitations, and indicate that impaired peroxisome degradation may not contribute to the pathology of MFF-deficiency. However, degradation of elongated peroxisomes in MFF-deficient cells may be slower than in control cells containing predominantly spherical peroxisomes, as tubules may need to shorten/fragment prior to removal by autophagy. Interestingly, a shortening of elongated peroxisomes was observed during amino acid starvation in HBSS, which was accompanied by alterations in peroxisomal marker proteins, e.g. the PMPs ACBD5 and PEX11β, which are required for membrane expansion and elongation. PEX11β mediates membrane deformation and elongation of the peroxisomal membrane (Delille et al. 2010; Opaliński et al. 2011), whereas ACBD5 has recently been shown to mediate membrane contact sites between peroxisomes and the ER by interacting with ER-resident VAP proteins (Costello et al. 2017b; Hua et al. 2017). Depletion of ACBD5 (or VAP) in MFF-deficient fibroblasts resulted in a shortening of elongated peroxisomes, likely due to disruption of the peroxisome-ER contact sites and reduced transfer of lipids from the ER to peroxisomes, which are required for peroxisomal membrane expansion (Costello et al. 2017b; Schrader et al. 2019). Our findings are in line with these previous observations and indicate that elongated peroxisomes in MFF-deficient cells are not fully static, but still dynamic under certain conditions. It is possible that a shortening/fragmentation of elongated peroxisomes under conditions of amino acid starvation facilitates their subsequent removal by autophagy.

Mitochondrial and peroxisomal dynamics are particularly important for brain development and function (Berger et al. 2016; Khacho and Slack 2018), likely explaining why MFF-deficient patients show primarily neurological defects. In contrast to the more prevalent neurological features in human patients with MFF-deficiency, mice without MFF die of heart failure at week 13, as a result of severe cardiomyopathy, which is likely based on mitochondrial alterations (Chen et al. 2015). However, removal of MFF exacerbated neuronal loss, astrogliosis and neuroinflammation in a Huntington's disease mouse model (Cha et al. 2018). Similar to patient fibroblasts, peroxisomes (and mitochondria) in MFF-deficient mouse embryonic fibroblasts were highly elongated (Chen et al. 2015). Interestingly, peroxisomal length was not substantially altered in MFF-deficient mouse cardiomyocytes (Chen et al. 2015). This strongly indicates that peroxisome morphology and division is affected in a cell type-specific manner.

We recently developed a mathematical model to explain and predict alterations in peroxisome morphology and dynamics in health and disease conditions (Castro et al. 2018). In this stochastic, population‐based modelling approach, each individual peroxisome consists of a spherical body with an optional cylindrical elongation. Peroxisome shape (i.e. the body radius and elongation length) are determined by (i) membrane lipid flow into the body (e.g., from the ER) (governed by rate *α* and lipid flow constant *γ*), (ii) elongation growth (governed by speed *v* and minimum body radius *r_min_*) and (iii) peroxisome division with a rate proportional to the elongation length (governed by rate *β* and minimum length *L*_min_). Peroxisome turnover is controlled by the peroxisome mean lifetime *τ*. We recently demonstrated that this model is applicable to a range of experimental and disease conditions, e.g. loss of PEX5 in Zellweger spectrum disorders (Castro et al. 2018). With wild-type parameters, peroxisomes in the model are typically high in number, with only a low percentage showing elongations, all of which are short (**Fig. 6A**). The morphological alterations of peroxisomes in MFF-deficient fibroblasts that we have observed experimentally are captured by changing only one parameter, namely by reducing the division rate *β* to almost zero (**Fig. 6B**). As the membrane lipid flow rate and elongation growth speed remain unchanged, this results in reduced numbers of peroxisomes with significantly longer membrane elongations (**Fig. 6D, E**). The observation that control fibroblasts display large numbers of small, spherical peroxisomes, but turn into few, extremely elongated organelles upon blocking of peroxisomal division, indicates that membrane lipid flow rate, elongation growth speed and division rate must be high in fibroblasts under normal conditions. In contrast, low membrane lipid flow rate or elongation speed in other cell types may result in a population of small peroxisomes and reduced numbers. This is reflected by depletion of ACBD5, which impacts on peroxisome-ER tethering and membrane expansion, resulting in shorter peroxisomes in MFF-deficient cells (Costello et al. 2017b). This morphological change can also be captured in the model by reducing the lipid flow rate *α* in addition to the division rate *β* (**Fig. 6C-E**). It is thus likely that peroxisome morphology is differently affected in various cell types in MFF-deficient patients. It should also be considered that environmental changes and related signalling events that trigger peroxisomal membrane expansion and division (e.g. metabolic alterations and certain stress conditions) can potentially promote the formation of hyper-elongated peroxisomes in formerly unaffected cell types and contribute to the pathophysiology of MFF-deficiency.

**Figure 6.**
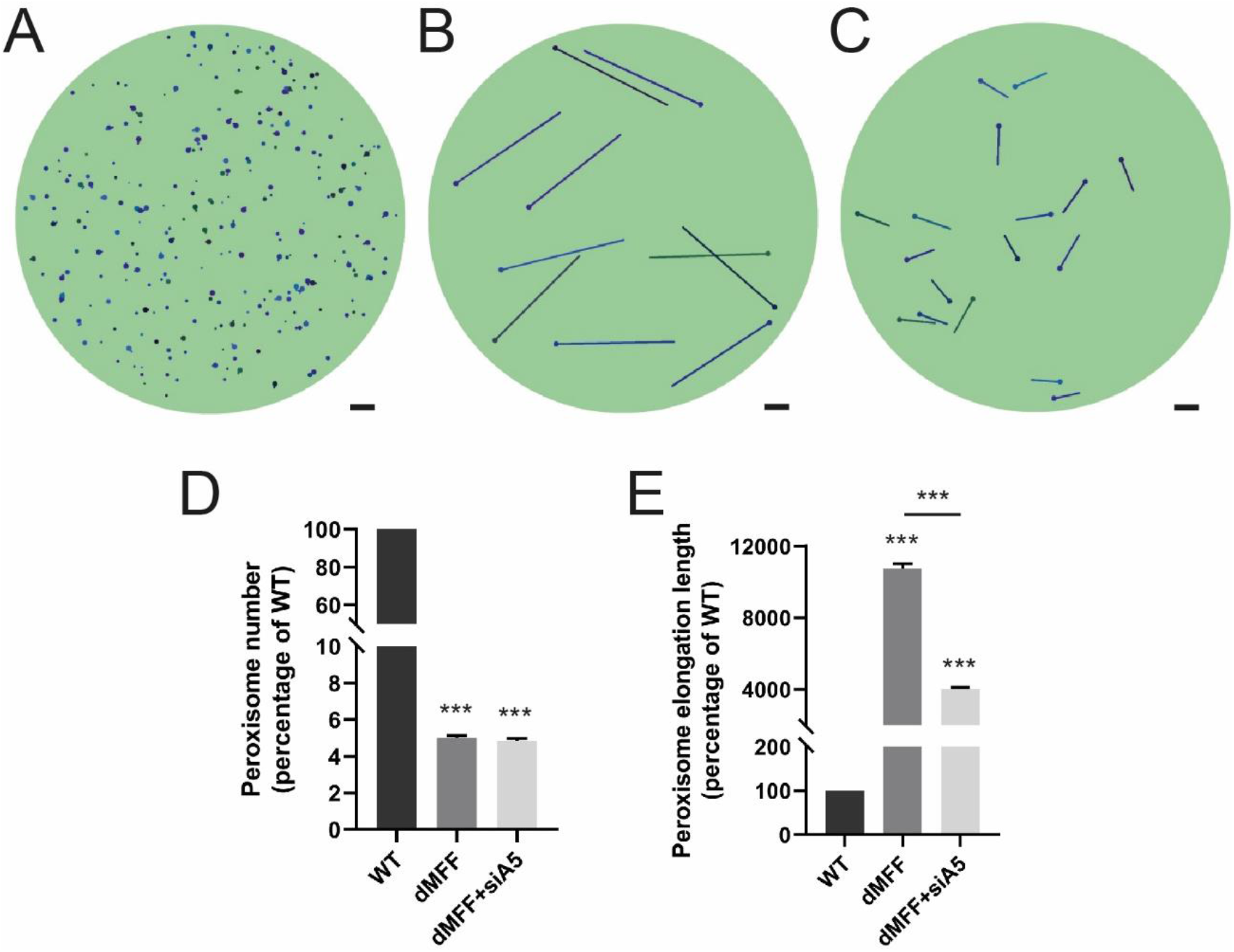
A mathematical model of peroxisome morphology and dynamics in wild-type and MFF-deficient patient fibroblasts. (**A**) Snapshot of model simulation for wild-type cells at *t* = 300 hours (α = 100 nm^2^/s, β = 2 × 10^−5^/nm/s, v = 0.3 nm/s, τ = 4 × 10^5^ s, γ = 2.5 × 10^−7^/nm^2^). (**B**) Snapshot of model simulation of MFF-deficient cells (dMFF) at *t* = 300 hours (α = 100 nm^2^/s, β = 2 × 10^−15^/nm/s, v = 0.3 nm/s, τ = 4 × 10^5^ s, γ = 2.5 × 10^−7^/nm^2^). (**C**) Snapshot of model simulation of MFF-deficient cells with reduced lipid flow to simulate silencing of ACBD5 (siA5) at *t* = 300 hours (α = 5 nm^2^/s, β = 2 × 10^−15^/nm/s, v = 0.3 nm/s, τ = 4 × 10^5^ s, γ = 2.5 × 10^−7^/nm^2^). (**D**) Average peroxisome number at *t* = 300 hours of simulations shown in **A**-**C**, represented as percentages relative to WT (n = 100). (**E**) Average non-zero peroxisome elongation length at *t* = 300 hours of simulations shown in **A**-**C**, represented as percentages relative to WT (n = 100). Scale bars, 1 μm.

## Abbreviations

ACOX1: acyl-CoA oxidase 1
PBD: peroxisome biogenesis disorder
PED: single peroxisomal enzyme deficiency
DRP1: dynamin-related protein 1
ER: endoplasmic reticulum
FIS1: mitochondrial fission 1 protein
MFF: Mitochondrial fission factor
ROS: reactive oxygen species
PTS: peroxisome targeting signal
VLCFA: very-long-chain fatty acid

## 5. Additional Information

### 5.1. Acknowledgements

We would like to thank F.S. Alkuraya, King Faisal Specialist Hospital and Research Center, Riyadh, Saudi Arabia for providing patient skin fibroblasts, and M. Schuster for support with FRAP experiments. This work was supported by the Biotechnology and Biological Sciences Research Council (BBSRC) (BB/N01541X/1, BB/R016844/1; to M.S.), the European Union’s Horizon 2020 research and innovation programme under the Marie Skłodowska-Curie grant agreement No 812968 PERICO (to M.S., M.F.), and the Research Foundation – Flanders (G095315N; to M.F.). M.I. was supported by the German Research Foundation (DFG Grant 397476530) and MEAMEDMA Anschubförderung, Medical Faculty Mannheim, University of Heidelberg. Y.W. was supported by the German Research Foundation (DFG Grant D10043030, SFB 1134 – Functional Ensembles). D.M.R. gratefully acknowledges financial support from the Medical Research Council (MR/P022405/1) and the Wellcome Trust Institutional Strategic Support Award (WT105618MA). C.L. was supported by the KU Leuven (PDM/18/188) and the Research Foundation – Flanders (1213620N). Support grants to J.P. were provided by Zellweger UK, The Sidney Perry Foundation and The Devon Educational Trust.

The research data supporting this publication are provided within this paper and as supplementary information.

The authors declare no competing financial interests.

### 5.2. Author Contributions

JP, RC, TS, LG, SF, CL, YW, CH, MI, DR, MS performed experiments and analysed data. MS, SF, PF, MF, MI conceived the project and analysed data. JP, MS wrote the manuscript. All authors contributed to methods.

---

**Table S1.**
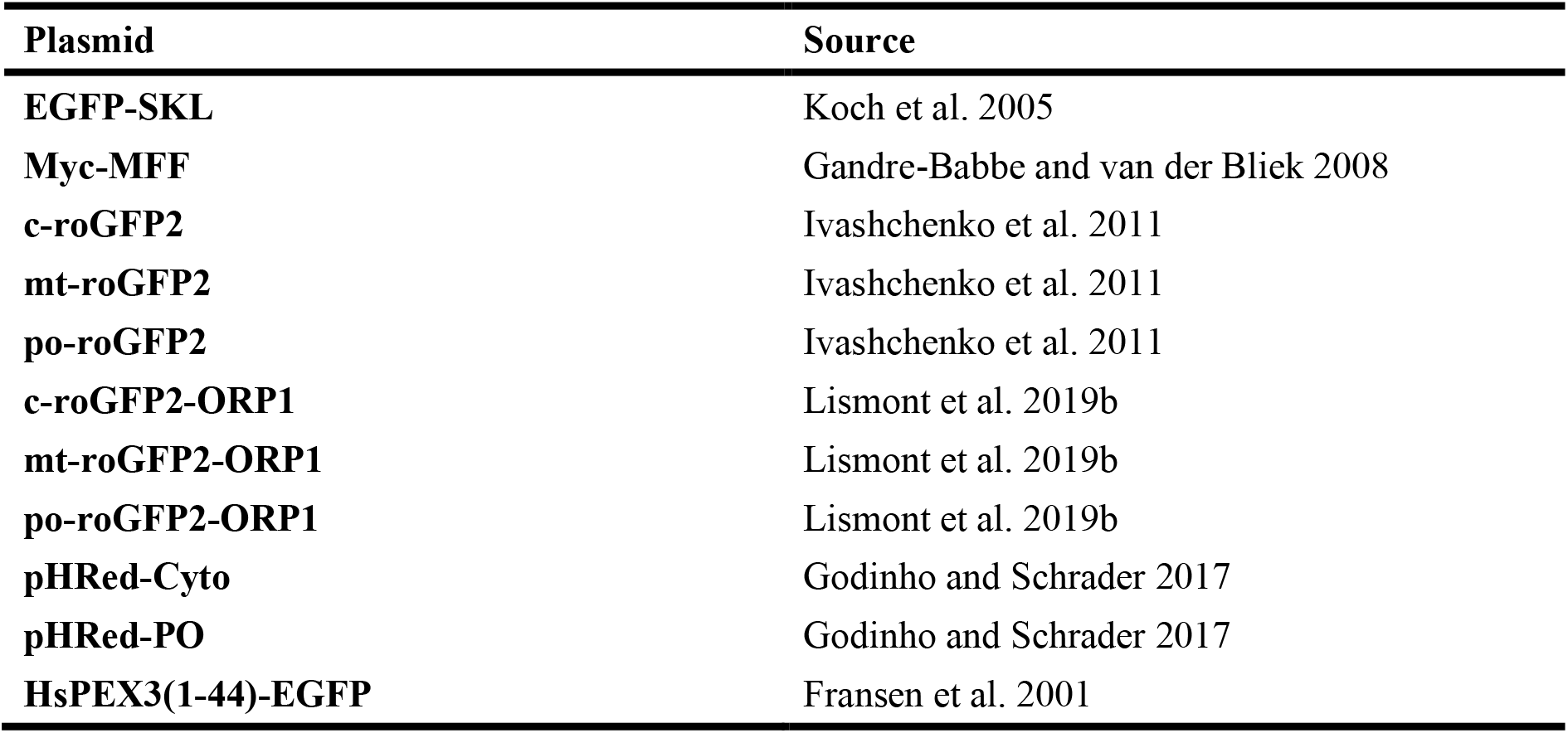
Plasmids used in this study

**Table S2.** Primary and secondary antibodies used in this study

| Antibody | Type | Dilution |  | Source |
| --- | --- | --- | --- | --- |
|  |  | IMF | WB |  |
| ACBD5 | pc rb |  | 1:1000 | Proteintech (21080-1-AP) |
| ACOX1 | pc rb | - | 1:1000 | Proteintech (10957-1-AP)<br>or gift from T. Hashimoto, Japan |
| ATP synthase | mc ms | 1:500 | - | Abcam (ab14730) |
| α-Tubulin | mc ms | - | 1:1000 | Sigma (T9026) |
| Catalase | pc ms | 1:150 | - | Abcam (ab88650) |
| Catalase | mc rb | - | 1:250 | Abcam (ab179843) |
| GAPDH | pc rb | - | 1:5000 | ProSci (3783) |
| Myc | mc ms | 1:200 | - | Santa Cruz Biotechnology, Inc (9E10) |
| PEX5 | pc rb | - | 1:750 | Sigma (HPA039259) |
| PEX11β | mc rb | | 1:1000 | Abcam (ab181066) |
| PEX14 | pc rb | 1:1400 | 1:4000 | D. Crane, Griffith University, Brisbane, Australia |
| PMP70 | pc rb | 1:100 | - | A. Völkl, University of Heidelberg, Heidelberg, Germany |
| PMP70 | mc ms | 1:500 | - | Sigma (SAB4200181) |
| **Thiolase** | pc rb | - | 1:2000 | Atlas antibodies (HPA007244) |
| **Alexa Fluor 488** | dk anti-ms | 1:500 | - | ThermoFisher Scientific (A21202) |
| **Alexa Fluor 488** | dk anti-rb | 1:500 | - | ThermoFisher Scientific (A21206) |
| **Alexa Fluor 594** | dk anti-ms | 1:500 | - | ThermoFisher Scientific (A21203) |
| **Alexa Fluor 594** | dk anti-rb | 1:500 | - | ThermoFisher Scientific (A21207) |
| **HRP IgG** | gt anti-ms | - | 1:10000 | Bio-Rad (170-6516) |
| **HRP IgG** | gt anti-rb | - | 1:10000 | Bio-Rad (172-1013) |
| **IRDye 800 CW** | gt anti-rb | - | 1:12500 | Westburg |

Abbreviations: IMF, immunofluorescence; WB, Western blot; pc, polyclonal; mc, monoclonal; ms, mouse; rb, rabbit; gt, goat; dk, donkey; HRP, horseradish peroxidase.

**Suppl. Figure S1.**
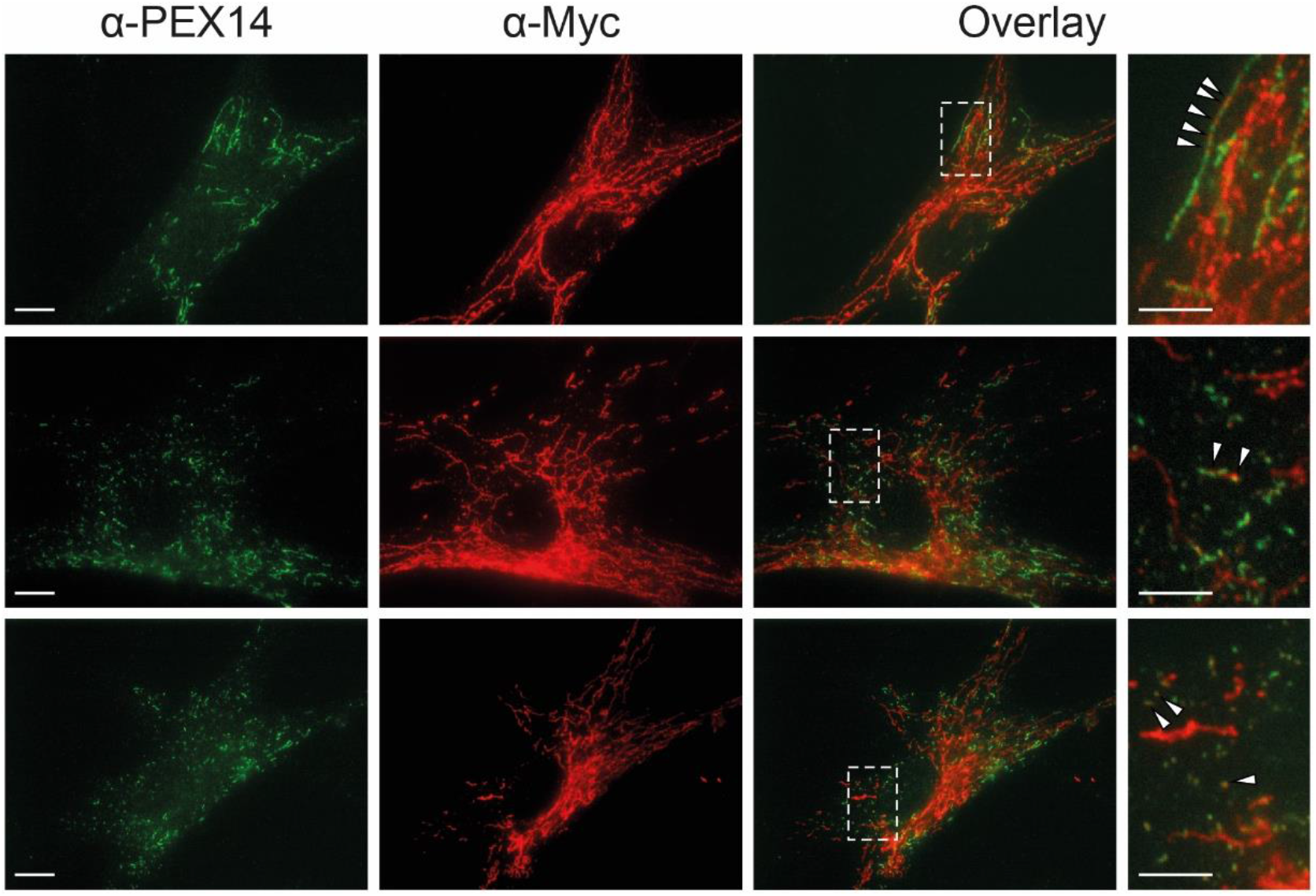
Re-introduction of MFF in MFF-deficient patient fibroblasts. MFF-deficient patient fibroblasts [mutation Q64* (Shamseldin et al. 2012)] were transfected with Myc-MFF using microporation. Cells were processed for immunofluorescence microscopy 2-3 hours after transfection using antibodies directed to the Myc-tag and PEX14, a peroxisomal membrane marker. Note the localisation of MFF in spots on elongated peroxisomes (upper panel; arrowheads), the appearance of shorter peroxisomes due to peroxisome division (middle panel), and the restoration of the normal, spherical peroxisome morphology (lower panel). Higher magnification of boxed regions is shown. Scale bars, 10 μm; magnification, 5 μm.

**Suppl. Figure S2.**
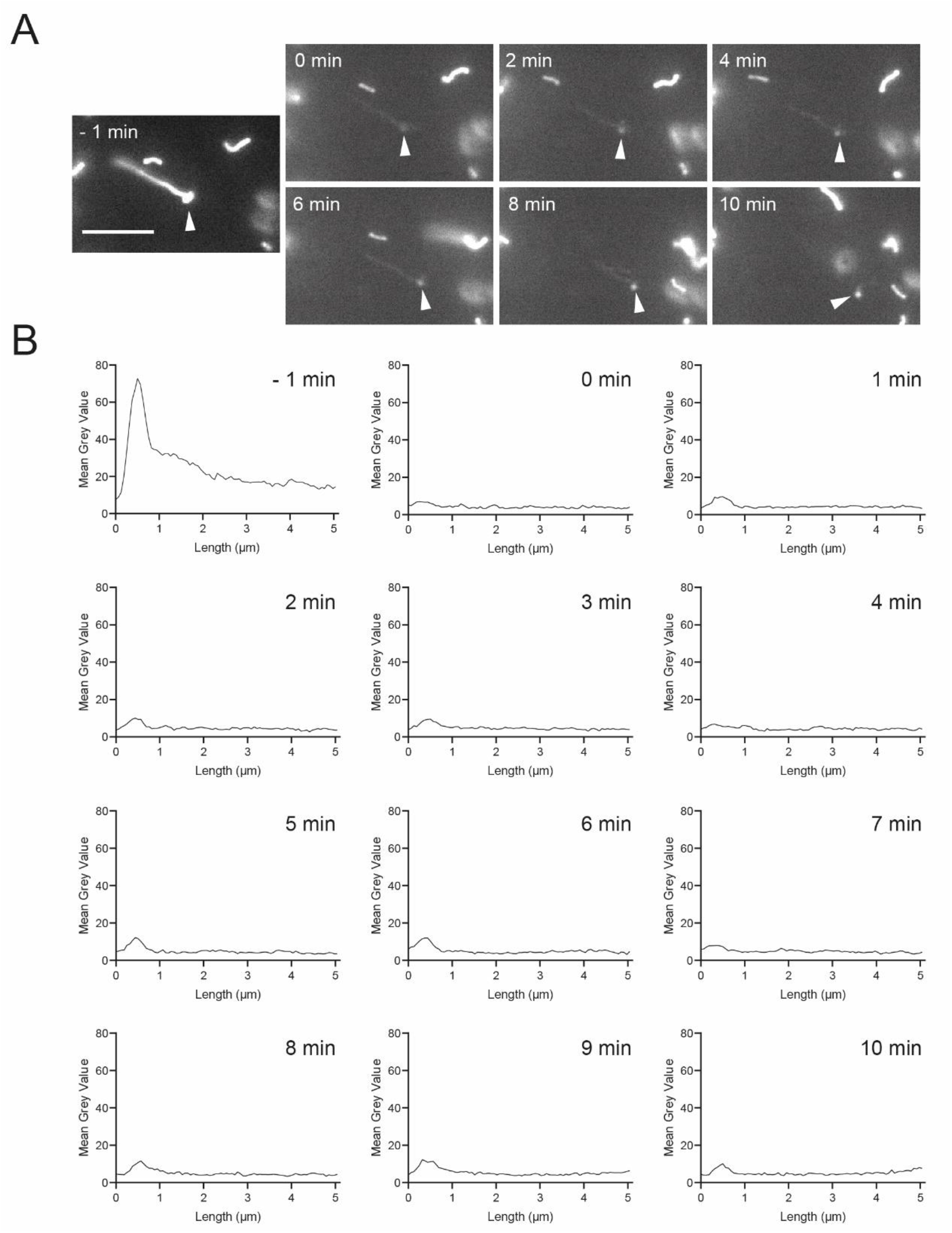
The peroxisomal body is import-competent. MFF-deficient fibroblasts were transfected with GFP-SKL and grown on 3.5-cm glass bottom dishes. Photo-bleaching experiments were performed after 24-48 hours using a Visitron 2D FRAP system. The entire organelle (peroxisome body and short tubule) was photo-bleached (0 min) and recovery of GFP-SKL fluorescence monitored over a period of 10 minutes (**A**). Note that GFP-SKL fluorescence was observed in the peroxisome body (arrowheads), but not in the peroxisome tubule, indicating that recovery is due to import of GFP-SKL into the peroxisome body. (**B**) Quantification of fluorescence intensity. Data are presented at the mean grey value for each increment along the length of the peroxisome. Scale bar, 5 μm.

**Suppl. Figure S3.**
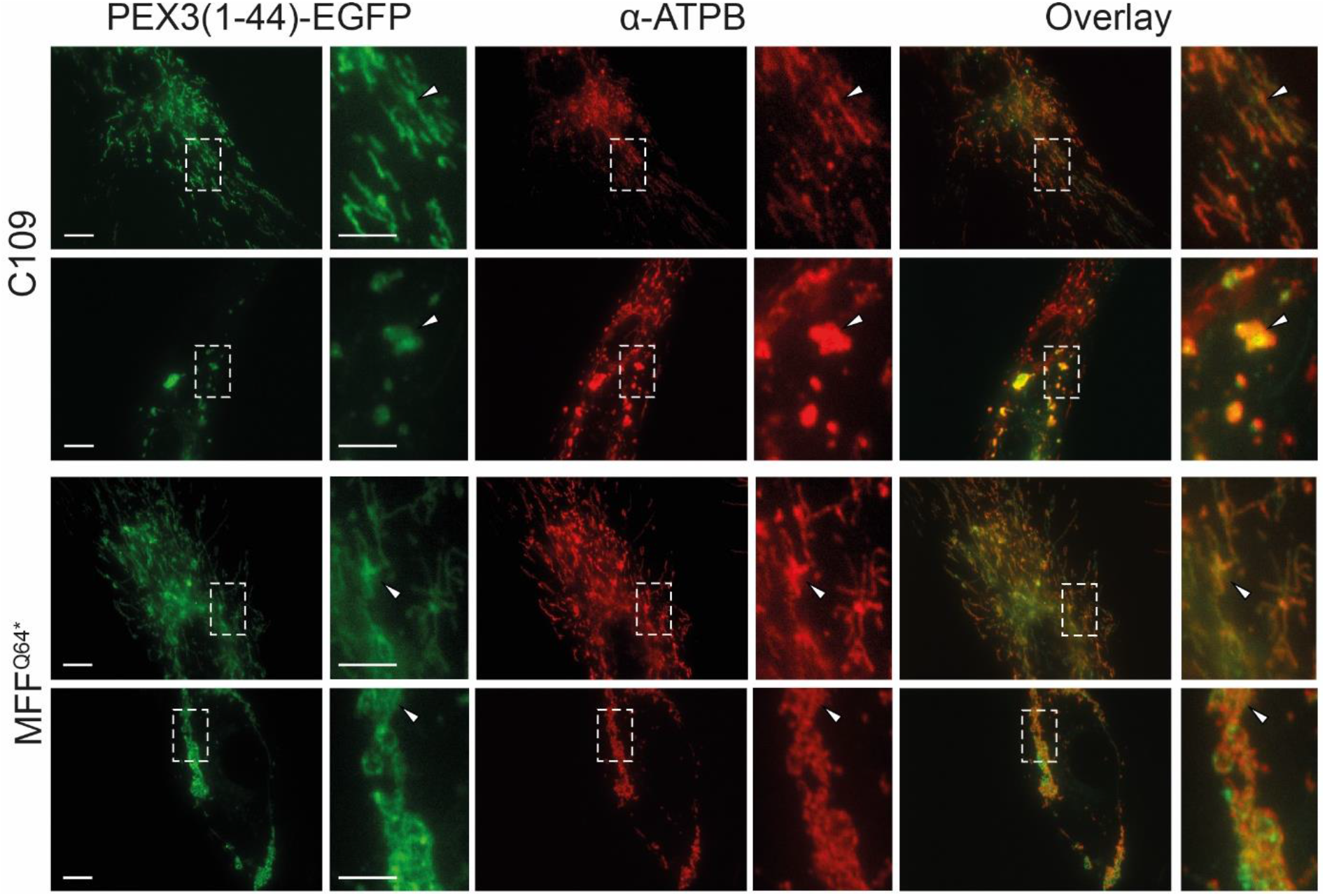
HsPEX3(1-44)-EGFP is targeted to mitochondria when peroxisomes are lost. Human control (C109) or MFF-deficient (MFF^Q64*^) fibroblasts were transfected with a plasmid coding for *Hs*PEX3(1-44)-EGFP to induce peroxisome degradation and processed for immunofluorescence after 24 and 48 hours using antibody against mitochondrial ATP synthase (ATPB). Note the mistargeting of *Hs*PEX3(1-44)-EGFP to mitochondria (arrowheads). Furthermore, mitochondrial morphology is altered including fragmentation and clustering. Scale bars, 10 μm, magnification, 2 μm.

**Suppl. Figure S4.**
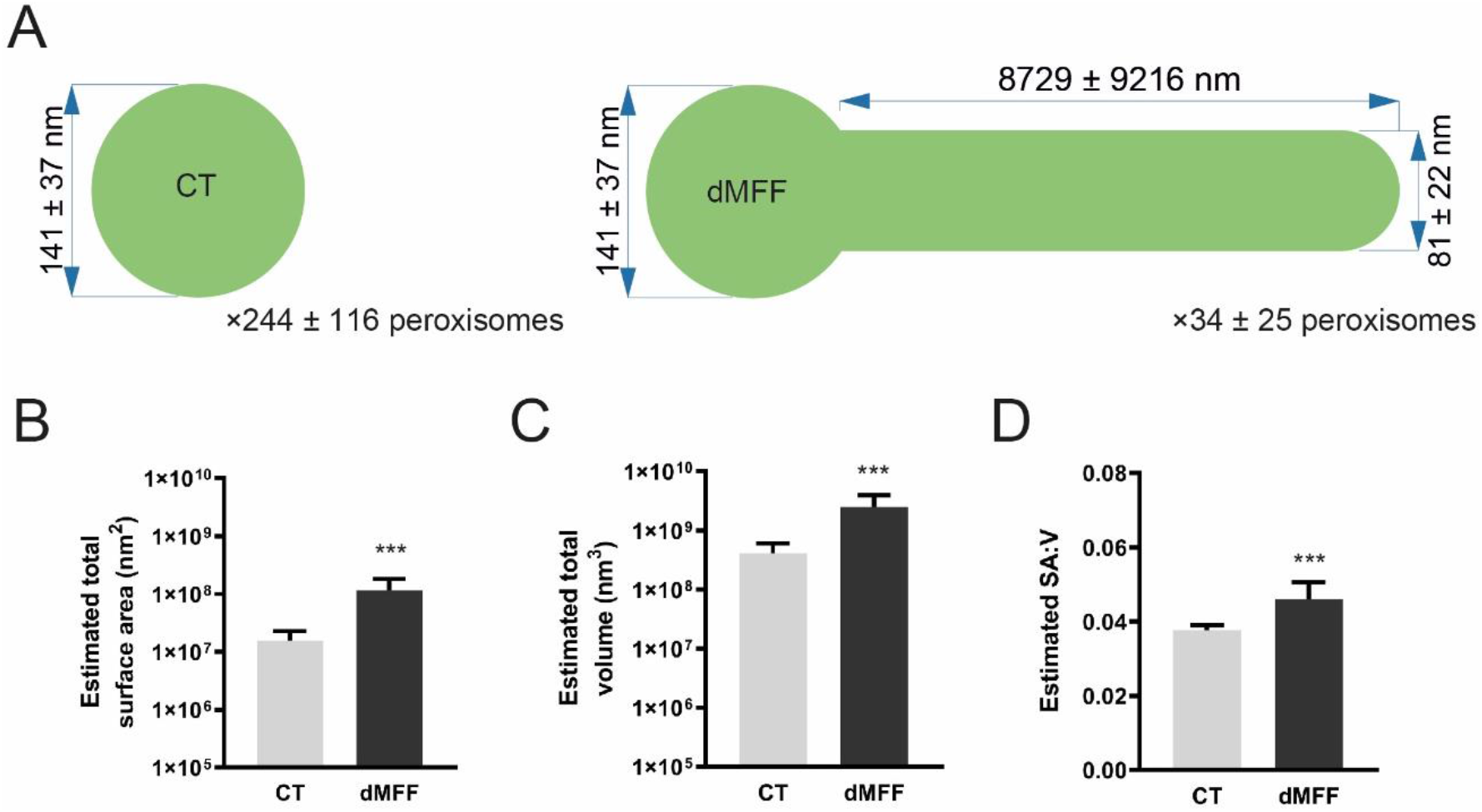
Calculations of peroxisomal surface area, volume, and surface area to volume ratio. (**A**) Values used for calculations (mean ± SD). For the non-elongated control (CT) body diameter, the value obtained from measurement of peroxisomal bodies in MFF^Q64*^ was used. (**B**) Estimated total peroxisomal surface area in control (CT) and MFF-deficient (dMFF) fibroblasts, based on an average of a computer-generated population of peroxisomes using values taken from the distributions shown in **A**. (**C**) Estimated total peroxisomal volume, and (**D**) estimated surface area to volume ratio (SA:V). Error bars show the mean + SD for 10,000 generated peroxisome populations. ***, *p* < 0.001; two-tailed, unpaired t test.

